# IMPDH2 filaments protect from neurodegeneration in AMPD2 deficiency

**DOI:** 10.1101/2024.01.20.576443

**Authors:** Marco Flores-Mendez, Laura Ohl, Thomas Roule, Yijing Zhou, Jesus A Tintos-Hernández, Kelsey Walsh, Xilma R Ortiz-González, Naiara Akizu

## Abstract

Metabolic dysregulation is one of the most common causes of pediatric neurodegenerative disorders. However, how the disruption of ubiquitous and essential metabolic pathways predominantly affect neural tissue remains unclear. Here we use mouse models of AMPD2 deficiency to study cellular and molecular mechanisms that lead to selective neuronal vulnerability to purine metabolism imbalance. We show that AMPD deficiency in mice primarily leads to hippocampal dentate gyrus degeneration despite causing a generalized reduction of brain GTP levels. Remarkably, we found that neurodegeneration resistant regions accumulate micron sized filaments of IMPDH2, the rate limiting enzyme in GTP synthesis. In contrast, IMPDH2 filaments are barely detectable in the hippocampal dentate gyrus, which shows a progressive neuroinflammation and neurodegeneration. Furthermore, using a human AMPD2 deficient neural cell culture model, we show that blocking IMPDH2 polymerization with a dominant negative *IMPDH2* variant, impairs AMPD2 deficient neural progenitor growth. Together, our findings suggest that IMPDH2 polymerization prevents detrimental GTP deprivation in neurons with available GTP precursor molecules, providing resistance to neurodegeneration. Our findings open the possibility of exploring the involvement of IMPDH2 assembly as a therapeutic intervention for neurodegeneration.

## Introduction

Protein aggregation is a hallmark of age-related neurodegenerative disorders that leads to a progressive loss of vulnerable neurons(Taylor *et al*, 2002). However, neurodegeneration can also occur in the absence of protein aggregation. Metabolic dysregulation, for example, is a common cause of neurodegeneration, especially in pediatric populations(Pierre, 2013). A significant proportion of inherited metabolic disorders show progressive decline of neurologic, cognitive and motor function as central signs of neurodegeneration(Saudubray & Garcia-Cazorla, 2018; Wong, 1997). In these conditions, neurodegeneration is associated with the accumulation of toxic metabolites, depletion of key metabolites, or bioenergetic defects.

Disruption of purine nucleotide metabolism underlies the pathogenesis of a large group of inherited metabolic disorders caused by mutations in key enzymes of the purine nucleotide synthesis pathway (Hartman & Buchanan, 1959). Purine nucleotides (i.e. ATP and GTP) supply cells with energy, constitute building blocks of DNA and RNA, and participate in intra- and inter-cellular signaling, therefore supporting cellular proliferation, survival and metabolic needs. To meet increased purine demands, proliferative cells synthetize purine nucleotides *de novo* in serial steps that convert phosphoribosyl pyrophosphate (PRPP) to inosine monophosphate (IMP) (Hartman & Buchanan, 1959; Lane & Fan, 2015; Watts, 1983). IMP is then used as common intermediate for *de novo* synthesis and interconversion of adenine and guanine nucleotides (i.e. ATP, ADP, AMP and GTP, GDP and GMP). In contrast, postmitotic cells favor the salvage pathway which recycles free purine bases into purine nucleotides (Hartman & Buchanan, 1959; Lane & Fan, 2015; Watts, 1983). The two pathways involve a series of sequential and often bidirectional reactions precisely coordinated by key metabolic enzymes.

The assembly of purine metabolism enzymes into micron-sized structures is emerging as a mechanism to fine tune and subcellularly compartmentalize purine nucleotide biosynthesis. For example, upon purine nucleotide starvation, up to 9 enzymes within the *de novo* purine biosynthetic pathway synchronize their function by assembling into dynamic, microscopically visible, intracellular metabolons known as purinosomes (An *et al*, 2008). Purinosomes crosstalk with mitochondria to power *de novo* purine biosynthesis when local purine nucleotide demands increase (French *et al*, 2016; Pareek *et al*, 2020). Another reaction catalyzed by the rate limiting enzymes of *de novo* guanine nucleotide biosynthesis pathway, IMPDH1 and IMPDH2, also involves supramolecular assemblies. Under physiological conditions that demand the expansion of guanine nucleotide pools, IMPDH1 and IMPDH2 assemble into large filaments (Gunter *et al*, 2008; Ji *et al*, 2006; Juda *et al*, 2014). The effect of these large filaments on IMPDH1/2 function is complex, but recent evidence suggest that they make the enzymes more resistant to allosteric inhibition by GTP, thus allowing to increase of guanine nucleotide pools even when the cellular supplies are high (Buey *et al*, 2015; Johnson & Kollman, 2020). Nevertheless, the implication of these macromolecular assemblies in physiological and pathological conditions remains poorly understood.

The nervous system is particularly vulnerable to imbalances of purine nucleotide and nucleoside pools given their additional functions as neuromodulators, neurotransmitters and secondary messengers (Badimon *et al*, 2020; Pascual *et al*, 2005). Accordingly, several inherited purine nucleotide metabolism disorders show neurologic manifestations (Camici *et al*, 2010). Motor and cognitive disability and self-injurious behavior are hallmarks of Lesch-Nyhan syndrome (LNS) caused by mutations in *Hypoxanthine Guanine Phosphoribosyl Transferase* (*HGPRT1*), a key enzyme in the purine nucleotide salvage pathway. HGPRT1 deficiency impairs the recycling of purine bases into purine nucleotides and leads to an overproduction of uric acid. However, treatments to reduce uric acid accumulation do not improve neurological phenotypes, which suggest an alternative neuropathogenic mechanism (Fu *et al*, 2015; Lesch & Nyhan, 1964). Likewise, overproduction of uric acid is a hallmark of PRPS1 superactivity disorder, characterized by gout which occasionally co-occurs with sensorineural deafness, cognitive deficits and hypotonia (Becker *et al*, 1988; Sperling *et al*, 1972). In contrast, loss of function mutations in *PRPS1*, which cause a spectrum of diseases with clinical features of diverse severity, including neurodegeneration, likely results from depletion of purine biosynthesis as indicated by reduced levels of purines and uric acid in urine and plasma of patients (Arts *et al*, 1993; de Brouwer *et al*, 2007; Kim *et al*, 2007; Synofzik *et al*, 2014). Furthermore, heterozygous mutations in *IMPDH2* have recently been associated with neurodevelopmental disorders and mutations in *IMPDH1* lead to common retinal degenerative disorders (i.e. retinitis pigmentosa and leber congenital amaurosis) (Bowne *et al*, 2002; Kennan *et al*, 2002). Mutations in *IMPDH1/2* alter the conformation of their filamentous assemblies, potentially leading to a dysregulation of the guanine biosynthetic pathway as a disease mechanism (Buey *et al*., 2015; O’Neill *et al*, 2023). While metabolic consequences of mutations in purine nucleotide enzymes have been extensively investigated, they usually do not explain the pathogenesis of neurologic manifestations. Furthermore, disease specific variability in clinical manifestations suggests tissue specific regulatory mechanism and vulnerabilities that remain unclear.

Inactivating mutations in *AMPD2* cause Pontocerebellar Hypoplasia type 9 (PCH9) (Akizu *et al*, 2013; Kortum *et al*, 2018). Pontocerebellar hypoplasias are a group of rare monogenic disorders characterized by a reduced volume of the brainstem and cerebellum with variable involvement of other brain structures, often caused by fetal onset neurodegeneration. Patients usually present developmental delay, feeding problems and motor abnormalities, with the most severe cases showing regression and early death due to respiratory problems (van Dijk *et al*, 2018). Current classification comprises 17 types of PCH which are distinguished by the genetic diagnosis, brain imaging and clinical features (Zakaria *et al*, 2023). Functional annotation of PCH associated genes suggest impaired protein synthesis, RNA or energy metabolism as the underlaying pathogenic mechanism (van Dijk *et al*., 2018). PCH9 is clinically distinguishable from the other PCH types by the ‘figure 8’ shape of the midbrain in axial brain images (Akizu *et al*., 2013). Additional clinical features of PCH9 include, global developmental delay, postnatal microcephaly, atrophy of the cerebral cortex and thinner corpus callosum (Akizu *et al*., 2013; Kortum *et al*., 2018). As exception, patients carrying a homozygous null mutation that affects only one of the *AMPD2* isoforms show intact brain structure but early onset upper motoneuron degenerative disorder classified as hereditary spastic paraplegia 63 (HSP63) (Novarino *et al*, 2014).

*AMPD2* is one of three mammalian adenosine monophosphate (AMP) deaminase paralogs involved in the conversion of AMP to inosine monophosphate (IMP), a key enzymatic step for guanine nucleotide synthesis (i.e. GTP) within the purine metabolism pathway. While *AMPD2* is ubiquitously expressed, the other two paralogues, *AMPD1* and *AMPD3* are restricted to specific cell types or lowly expressed. *AMPD1* is predominantly expressed in the muscle, and homozygous mutations result in exercise-stress-induced muscle weakness and cramping (Fishbein *et al*, 1978), whereas *AMPD3* is predominantly expressed in erythrocytes, and homozygous mutations result in asymptomatic erythrocytic AMP accumulation (Zydowo *et al*, 1989). We previously showed that mutations in AMPD2 block the *de novo* and salvage production of GTP in PCH9 patient derived neural progenitor cell (NPC) cultures (Akizu *et al*., 2013). Partial loss of brain GTP levels and neurodegeneration were recapitulated in AMPD deficient mice only upon the double deletion of *Ampd2* and *Ampd3*. Furthermore, AMPD deficiency in mice led to premature death before postnatal day 21 preceded by signs of selective hippocampal neurodegeneration. This difference with PCH9 patients, who show a predominant involvement of the midbrain and cerebellum, suggests species-specific vulnerabilities to selective neuronal degeneration by AMPD deficiency. However, mouse hippocampal neurodegeneration is a useful model to study cellular and molecular features that mediate vulnerability to AMPD deficiency.

Here, we show that selective neuronal vulnerability to AMPD deficiency is inversely correlated with buildup of micron-sized IMPDH2 filaments. Taking advantage of AMPD deficient mice we uncover a selective accumulation of IMPDH2 filaments in the hippocampal CA1-3 while the dentate gyrus shows signs of neuroinflammation and lack of IMPDH2 filaments. To assess the effect of IMPDH2 aggregation in neurodegeneration further, we generated a forebrain specific AMPD deficient mice that survive to adulthood. Longitudinal histological analysis revealed a severe neurodegeneration of the hippocampal dentate gyrus while CA1-3 regions were free of reactive microglia and did not degenerate. Remarkably, CA1-3 regions of the hippocampus persistently showed an accumulation of IMPDH2 filaments, while these were sparse and thin in the dentate gyrus. Finally, we demonstrate that PCH9 patient derived NPCs also accumulate IMPDH2 filaments and their disassembly compromises PCH9 NPC growth. Altogether our work suggests induction of IMPDH2 filament assembly as a potential therapeutic intervention for PCH9 associated neurodegeneration.

## Results

### AMPD deficiency leads to selective hippocampal neurodegeneration in mice

Unlike *AMPD2* deficiency in humans, which causes PCH9, *Ampd2* deficient mice are behaviorally and neuropathologically indistinguishable from their control littermates likely due to compensation by brain *Ampd3* expression(Akizu *et al*., 2013; Toyama *et al*, 2012). However, we previously showed that the homozygous deletion of both *Ampd3* and *Ampd2* (*Ampd2*^-/-^;*Ampd3*^-/-^, hereafter referred as dKO) leads to premature death at postnatal day 21 and signs of neurodegeneration, particularly affecting the hippocampus. To further determine the extent of neurodegeneration in dKO mice, we performed several morphological and histological analysis. As previously shown dKO mice were born at normal Mendelian ratio and survived up to postnatal day 21 (p21) showing weakness, motor difficulties and growth restrictions as indicated by the smaller body size than their control littermates (Fig 1A). Consistently, dKO mice also showed smaller brains at postnatal day 20 (p20) (Fig 1B). To assess if the smaller brain was the result of a global neurodegeneration or part of the growth impairment, we monitored the progression of brain/body weight over the first 20 days of life in dKO and control littermates. Although dKO mice showed smaller brain than their control littermates, this was proportional to the reduced body weight at all analyzed timepoints (Fig 1C). This data indicates that the smaller brain likely results from the overall impaired growth of the dKO mice rather than microcephaly due to neurodevelopmental deficits or generalized neurodegeneration.

**Figure 1:**
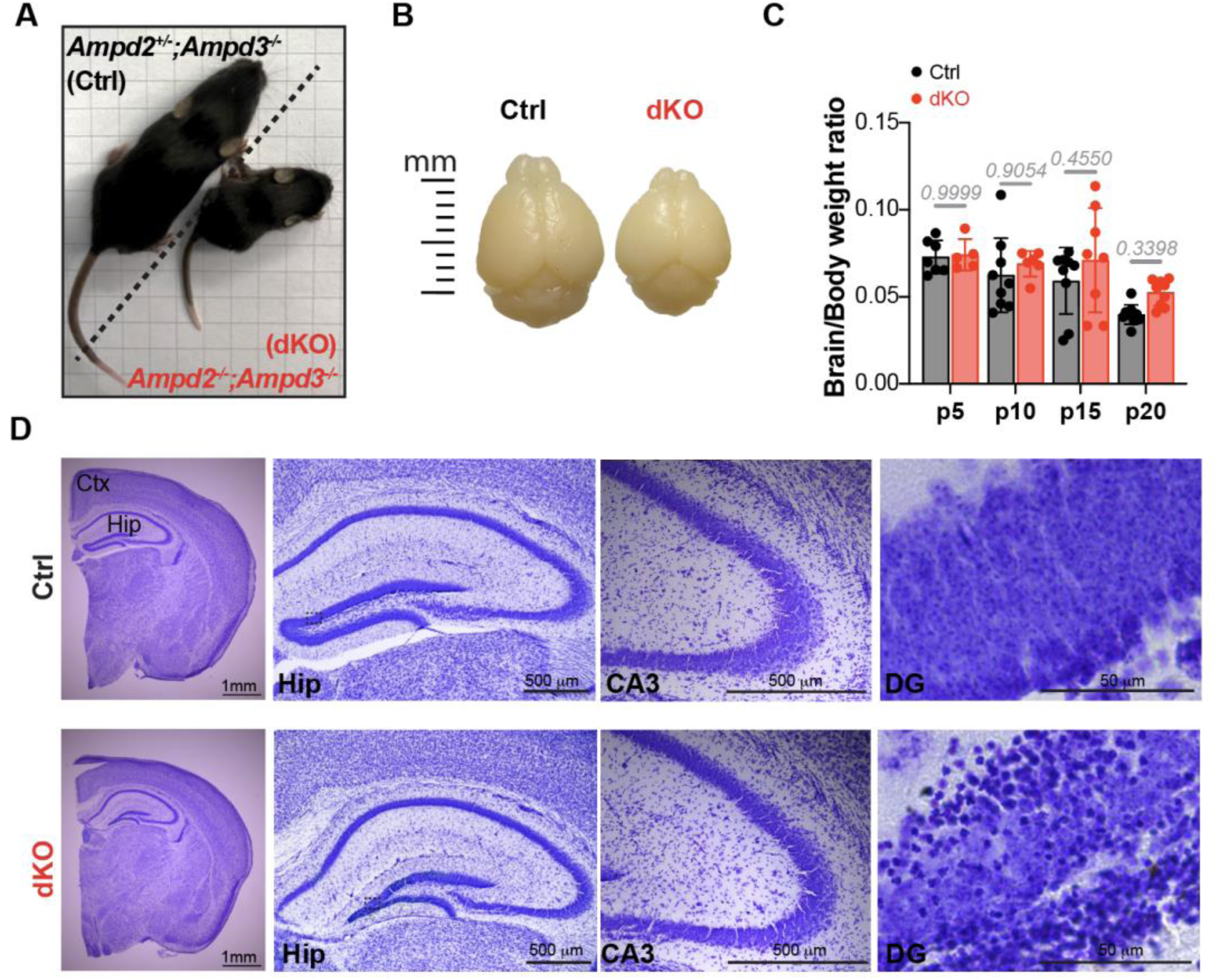
Ampd2 and Ampd3 double knock out (dKO) mice show selective hippocampal neurodegeneration. **(A)** Representative image of control (Ctrl) and double knockout (dKO) mice at postnatal day 20 (p20). **(B)** Brain images from Ctrl and dKO mice (p20) showing smaller size in dKO. **(C)** Brain and body weight ratio at different postnatal days (5, 10, 15 and 20 days). Bar graph shows mean ± SD of n=5-10 animals per genotype and time point represented by each dot in the graph. Significance was calculated with one way ANOVA with Sidak’s post hoc analysis for multiple comparison**. (D)** Nissl staining of brain coronal sections from Ctrl and dKO mice at p20 showing pyknotic cells in dKO hippocampus. Ctx=Cortex; Hip=Hippocampus; CA3=Cornu Ammonis; DG=Dentate Gyrus.

Nevertheless, since neurodegeneration is a hallmark of PCH9 and signs of neuronal death were previously observed in AMPD deficient mice (Akizu *et al*., 2013), we proceeded to analyze dKO mouse brains at histologic level. To this end, we collected dKO mice brains at the oldest age possible (p20). Histologic analysis showed structurally and histologically normal cerebellum and cerebrum in dKO mice (Fig 1D and S1A). However, in agreement with our previous work, dKO mice showed hippocampal deficiencies, particularly a thinner dentate gyrus with presence of pyknotic cells (Fig 1D), suggesting that AMPD deficiency in mice leads to selective hippocampal neurodegeneration.

### AMPD deficiency induces IMPHD2 aggregation in the hippocampus

Having established that AMPD deficiency in mice leads to a predominant neurodegeneration of the hippocampus, we next sought to determine causes of hippocampal vulnerability. AMPD2 deficiency blocks AMP to IMP conversion, which leads to a depletion of GTP worsened by the inhibition of the *de novo* purine biosynthetic pathway by adenine nucleotide accumulation (Akizu *et al*., 2013). Thus, we hypothesized that selective hippocampal vulnerability in dKO mice may result from a more severe purine nucleotide imbalance than in other brain regions. To test this hypothesis, we isolated hippocampi, cerebral cortices, and cerebella from dKO mice and control littermates at p20 and analyzed purine nucleotide levels by LC/MS. Results showed nearly intact ATP levels (Fig 2A), but an accumulation of AMP and ADP in all the dKO brain regions analyzed (Fig S2A). In contrast, IMP, which is the direct product of AMP deamination by AMPD enzymes, was among the most reduced nucleotide in all analyzed dKO brain areas (Fig 2B). Likewise, guanine nucleotides, which are produced from IMP by IMPDH2, were similarly reduced in dKO hippocampus, cortex and cerebellum (Fig 2C and S2B), indicating that nucleotide imbalance alone does not explain selective hippocampal vulnerability.

**Figure 2:**
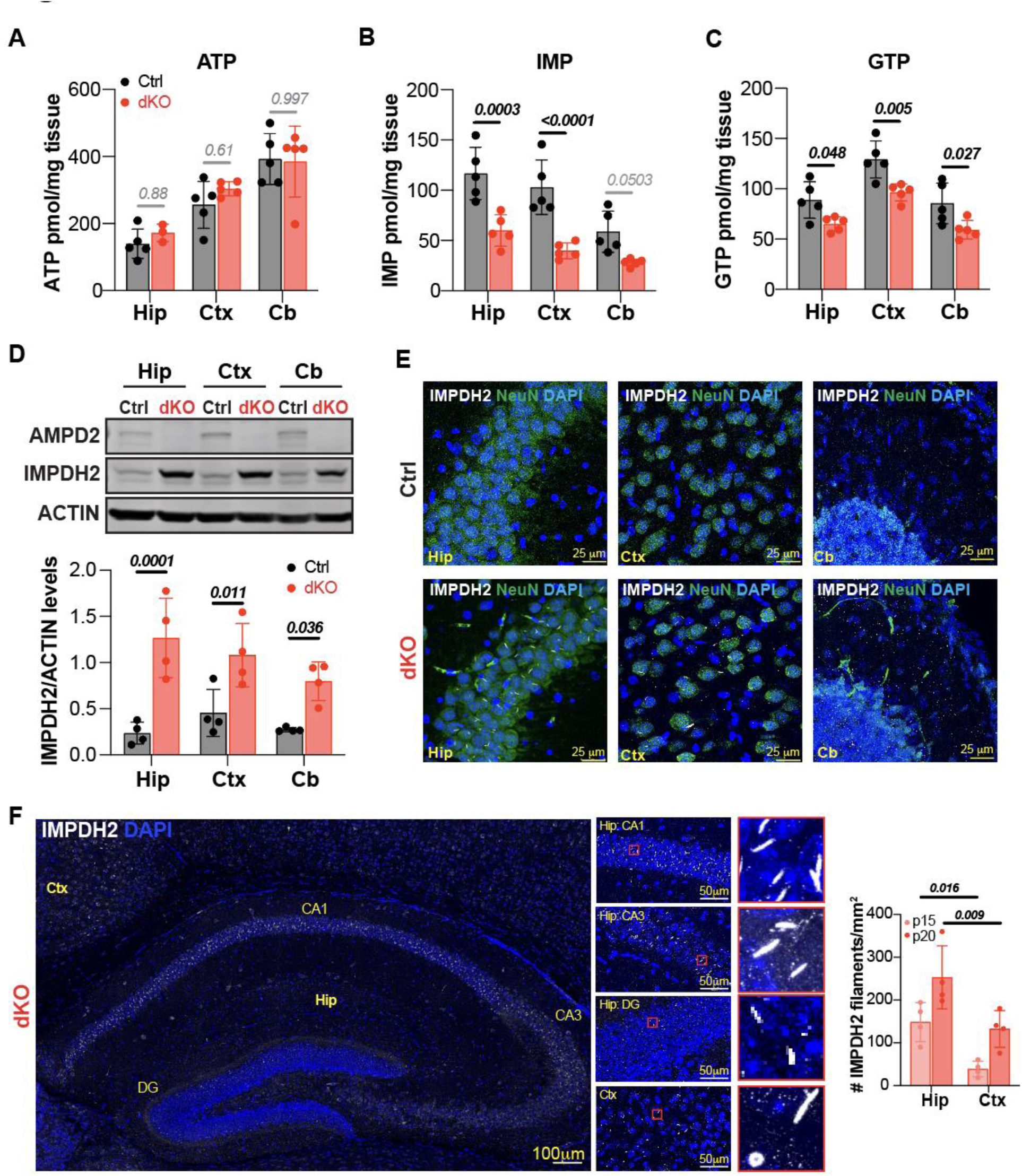
Hippocampal neurodegeneration is associated with IMPDH2 filament accumulation in dKO mice. **(A-C)** Nucleotide levels measured by LC/MS in Hippocampus (Hip), Cortex (Ctx) and Cerebellum (Cb) of control (Ctrl) and double knockout (dKO) mice at postnatal day 20 (p20) show similar ATP levels between dKO and control mice and reduction of IMP and GTP levels across all the brain regions of dKO mice compared to controls. Graphs show mean ± SD of n=5 animals per genotype. Significance was calculated with one way ANOVA and Sidak’s post hoc analysis for multiple comparison**. (D)** Representative western blot analysis showing absence of AMPD2 and upregulation of IMPDH2 in dKO compared to control mice at p20. ACTIN is shown as loading control. Graph shows mean ± SD of IMPDH2 band densitometry quantification relative to ACTIN in n=4 animals per genotype. Significance was calculated with one way ANOVA and Sidak’s post hoc analysis for multiple comparison**. (E)** Representative immunostainings of IMPDH2 and NEUN show IMPDH2 filaments in dKO brain hippocampus and cortex but not in controls. Cerebella are free of IMPDH2 filaments. **(F)** Representative immnunostaining of IMPDH2 in dKO mouse coronal sections showing abundant IMPDH2 filaments in Hippocampal (Hip) CA1-3 (Hip) regions. Right panels show magnified images. Graph shows mean ± SD of IMPDH2 filament density in the Hipocampus (Hip) and Cortex (Ctx) of p15 and p20 dKO mice. n=4 mice per genotype. Significance was calculated with one way ANOVA and Sidak’s post hoc analysis for multiple comparison. CA=Cornu Ammonis; DG=Dentate Gyrus.

To further validate dKO mice nucleotide deficits with an orthogonal approach, we took advantage of the regulation of IMPDH2 expression by guanine nucleotides. In yeast and mammalian cell cultures, depletion of guanine nucleotide pool triggers the overexpression of IMPDH2 and their homologues (Escobar-Henriques & Daignan-Fornier, 2001; Zimmermann *et al*, 1998). In agreement, Western Blot (WB) analyses showed that reduced guanine nucleotides in dKO mice brain regions are associated with an increase of IMPDH2 protein levels (Fig 2D). Furthermore, immunofluorescence stainings uncovered micron-sized filamentous IMPDH2 structures in dKO mice brain sections but not in controls (Fig 2E). Interestingly, despite the similarity in guanine nucleotide depletion and IMPDH2 overexpression of all analyzed brain regions, IMPDH2 filaments were most abundant in the hippocampus, particularly in CA1-3 regions, while only few were detected in the cortex and nearly none in the cerebellum (Fig 2E). Remarkably, IMP to GTP ratio, which plays an important role on IMPDH2 filament assembly (Johnson & Kollman, 2020), was also the largest in the hippocampus (Fig S2C). Most of these filaments had a rod shape with some arranged as a ring (Fig 2F). Despite their similarity with primary cilia, co-immunostaining with acetylated tubulin confirmed that IMPDH2 rods and rings are unrelated to primary cilia (Fig S2D). Instead, by ultrastructure analysis, we detected a perinuclear enrichment of IMPDH2 filaments often showing mitochondria in the proximity (Fig S2E). Overall, these data show that the predominant vulnerability of the hippocampus to GTP depletion is associated with a disproportionate IMPDH2 filament assembly.

### Brain region specific transcriptomes reveal apoptosis and neuroinflammation signatures in dKO mice hippocampus

To determine why the hippocampus is more vulnerable to GTP depletion and IMPDH2 assembly in dKO mice, we conducted a mRNA sequencing (RNAseq) analysis of the three brain regions exhibiting different degrees of neurodegeneration and IMPDH2 aggregation: the hippocampus showing neurodegeneration and IMPDH2 filaments, the cerebral cortex with apparently no neurodegeneration and low density of IMPDH2 filaments and the cerebellum with no degeneration or IMPDH2 filaments. We first analyzed the RNAseq data to identify differentially expressed genes (DEGs) between control and dKO mice in each brain region. Interestingly, the hippocampus exhibited the largest number of DEGs with 342 genes downregulated and 328 upregulated in dKO compared to control mice (Fig 3A-C). Functional annotation analysis of the three brain regions also showed a significant enrichment of ‘mTORC1 signaling’ pathway among downregulated DEGs, with 19 genes contributing to this category in the hippocampus, 6 in the cortex and 14 in the cerebellum (Fig 3D). Remarkably, depletion of guanine nucleotides is known to inhibit mTORC1 activity in order to compensate for the high nucleotide demand of mTORC1-stimulated ribosomal RNA synthesis (Hoxhaj *et al*, 2017). Therefore, the downregulation of mTORC1 signaling genes may be a consequence of the overall guanine nucleotide reduction in dKO mice. To test for this possibility and assess for brain region specific effects on mTORC1 inhibition, we extracted proteins from hippocampus, cortex and cerebellum of control and dKO mice and analyzed ribosomal protein S6 phosphorylation, as a proxy of mTORC1 signaling activity. Results confirmed mRNA-seq data and indicated that mTORC1 signaling inhibition is generalized in the three brain regions of dKO mice (Fig 3E), ruling out mTORC1 inhibition as the underlaying cause of the selective hippocampal neurodegeneration.

**Figure 3:**
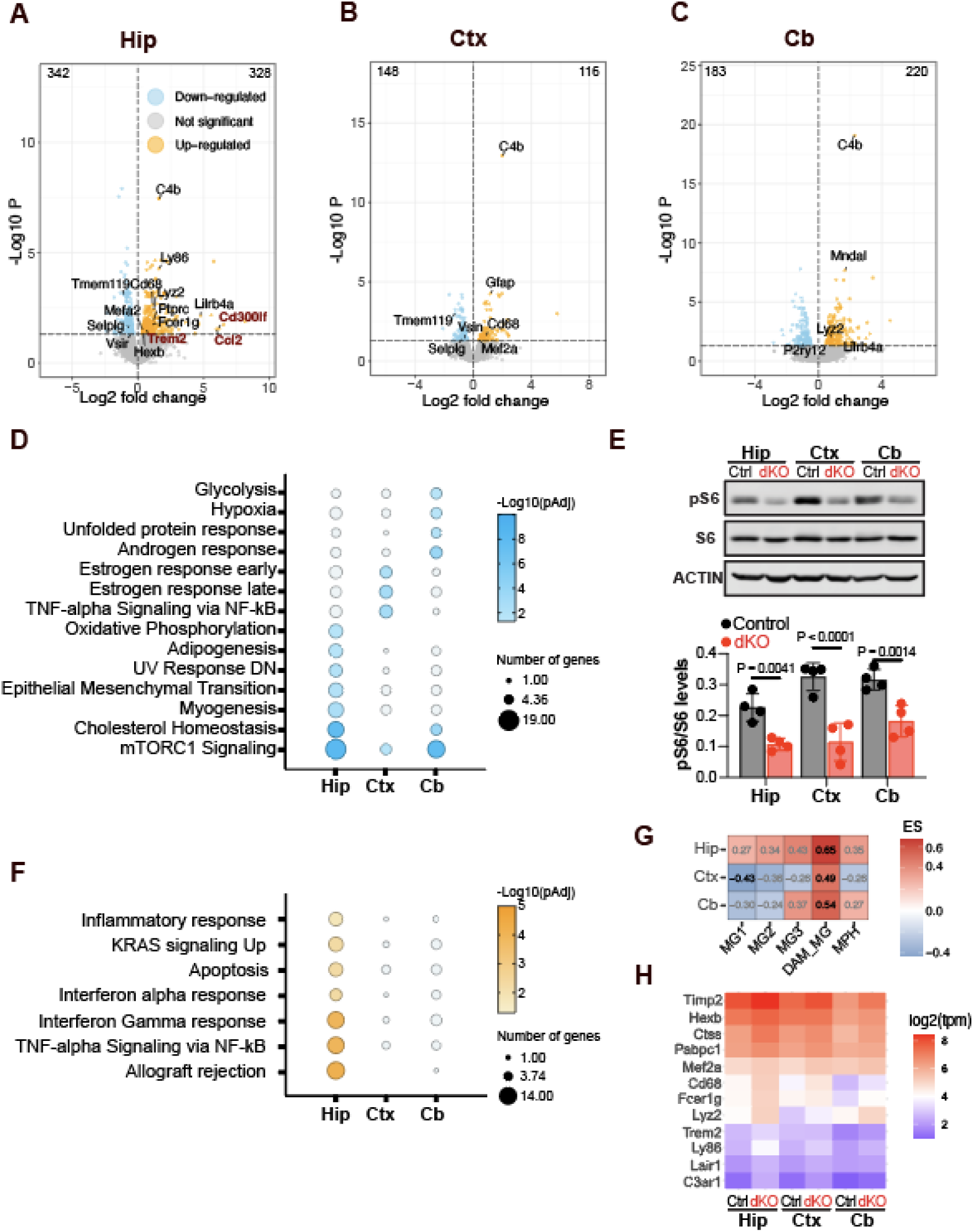
dKO mice brains show generalized mTORC1 signaling inhibition and hippocampal selective neurodegeneration and neuroinflammation transcriptomic signatures. **(A-C)** Volcano plots of differentially expressed genes (DEGs) in Hippocampus **(A)**, Cortex **(B)** and Cerebellum **(C)** of dKO vs Ctrl mice at postnatal day p20 (n=3 mice per genotype). Horizontal dashed lines indicate statistical significance cut off (p<0.05 or -log10>1.301). **(D)** Bubble plot showing functional annotation analysis of downregulated DEGs in dKO vs Ctrl Hippocampus (Hip), Cortex (Ctx) and Cerebellum (Cb). White circles are non-significant. **(E)** Representative western blot analysis showing reduction of pS6 in dKO Hippocampus (Hip), Cortex (Ctx) and Cerebellum (Cb), which indicates mTORC1 inhibition. ACTIN is shown as loading control. Graph shows mean ± SD of pS6 band densitometry quantification relative to total S6 in n=4 animals per genotype. Significance was calculated with one way ANOVA and Sidak’s post hoc analysis for multiple comparison**. (F)** Bubble plot showing functional annotation analysis of upregulated DEGs in dKO vs Ctrl Hippocampus, Cortex and Cerebellum. White circles are non-significant. **(H)** GSEA analysis of microglia gene signatures in in dKO vs Ctrl Hippocampus, Cortex and Cerebellum DEGs. **(G)** Heatmap of microglia gene markers expression (log2(TMP)) in dKO and Ctrl Hippocampus, Cortex and Cerebellum.

On the other hand, functional annotation analysis of upregulated DEGs revealed a significant enrichment of apoptosis associated genes only in the dKO hippocampus (Fig 3F). This data is in line with the selective hippocampal neurodegeneration we found by histological analysis. Moreover, only DEGs upregulated in the hippocampus were enriched for inflammatory response, interferon and TNF alpha response pathways, all associated with neuroinflammation (Fig 3F). To further confirm that hippocampal selective neurodegeneration is associated with an inflammatory response, we conducted a gene set enrichment analysis (GSEA) of DEGs in the three brain regions using as input microglia specific gene signatures curated from published single-cell RNA sequencing datasets (Keren-Shaul *et al*, 2017). Interestingly, the dKO hippocampus showed a greater enrichment of microglia associated genes than the cortex and cerebellum DEGs including a significant enrichment for disease associated microglia genes (Fig 3G). Consistently, the expression levels of several microglia marker genes were also the highest in the dKO hippocampus (Fig 3H). These data suggest that GTP depletion and IMPDH2 filament accumulation in the hippocampus is associated with microglia accumulation.

### Microglia accumulation is restricted to regions of the hippocampus with low density of IMPDH2 filaments

IMPDH2 is well known for its role in supporting metabolic needs of immune system, particularly during T-cell stimulation (Duong-Ly *et al*, 2018; Zimmermann *et al*., 1998). Furthermore, recent data suggest that IMPDH2 may be involved in brain microglia infiltration (Liao *et al*, 2017). Based on these data, we considered the possibility that IMPDH2 filaments in dKO hippocampus may result from IMPDH2 assembly in infiltrating or reactive microglia. To test this hypothesis, we co-immunostained brain sections with anti-CD68 microglia marker and anti-IMPDH2. Results confirmed an accumulation of microglia only in the dKO mice and predominantly in the hippocampus (Fig 4A). Surprisingly, microglia primarily accumulated in the DG (Fig 4A), the region of the hippocampus with the lowest density of IMPDH2 filaments (Fig 2F and Fig 4B). In contrast, CA1-3 region of the hippocampus, where IMPDH2 filaments mostly accumulate (Fig 2F), were nearly free of microglia (Fig 4A, B). Thus, our results demonstrated that IMPDH2 filaments do not accumulate in CD68+ microglia (Fig 4B). We then explored the possibility of IMPDH2 filament accumulation in astrocytes, which are also key regulators of neuroinflammation, but co-immunostaining with anti-GFAP confirmed absence of IMPDH2 filaments in astrocytes (Fig 4C). These data left neurons as the most plausible candidates to accumulate IMPDH2 filaments. Accordingly, anti-NeuN immunostaining showed co-localization of IMPDH2 filaments with NeuN+ hippocampal CA1-3 neurons (Fig 4C).

**Figure 4:**
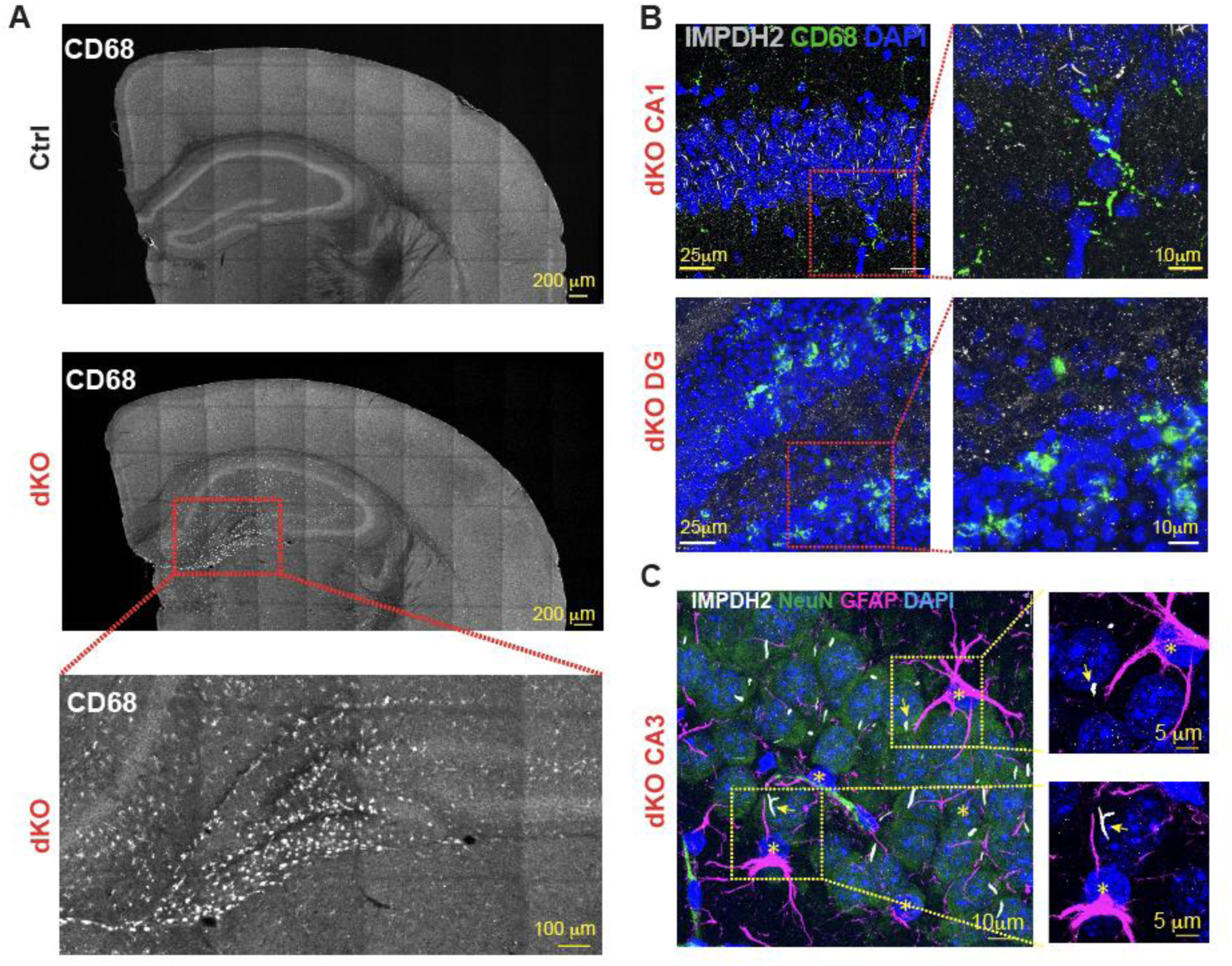
IMPDH2 filaments and microglia accumulation are inversely correlated in dKO hippocampus. **(A)** Representative images of coronal brain sections from Control (Ctrl) and double Knockout (dKO) mice at postnatal day 20 (p20) immunostained with anti-CD68 antibody. Inset magnification (red square) shows hippocampal DG regions with abundant CD68+ microglia in dKO mouse. **(B)** Immunostaining for IMPDH2 and CD68 in hippocampal regions (CA1 and DG) of dKO mice. Upper inset magnification (red square) shows hippocampal CA1 (upper panel) region with few CD68+ microglia which are free of IMPDH2 filaments. Lower inset magnification shows hippocampal DG (red square) where neurons without IMPDH2 filaments are surrounded by CD68+ microglia. **(C)** Representative images of hippocampal CA3 region co-immunostained with anti-IMPDH2, anti-NEUN neuronal marker and anti-GFAP astrocyte marker (GFAP) showing localization of IMPDH2 filaments predominantly in NEUN+ neurons.

### IMPDH2 filament accumulation in CA1-3 associates with resistance to neurodegeneration

The accumulation of reactive microglia is often associated with disease progression in neurodegeneration. Thus, we wondered whether neurons in the microglia rich DG of dKO mice were more prone to degenerate than neurons in microglia free regions (i.e. CA1-3). However, the premature death of dKO at p21 precluded us from assessing neurodegeneration further. To overcome this challenge, we sought to generate an AMPD deficient mouse model that survived longer by deleting AMPD activity only in the forebrain. To this end, we created a conditional *Ampd3* allele by removing the LacZ trapping cassette from the *Ampd3* knockout-first allele and subsequently breed it into *Ampd2* knock out background and Emx1:Cre knock in mice. This strategy produced a forebrain specific AMPD deficient mouse hereafter named cdKO (Fig S3A). The upregulation of IMPDH2 in cerebral cortex and hippocampus observed by WB analysis confirmed the deletion of AMPD activity in forebrain regions (cortex and hippocampus) (Fig S3B). In contrast, IMPDH2 levels in the cdKO cerebellum were comparable to control mice (Fig S3B) as predicted by the lack of Emx1:cre expression in the cerebellum (and therefore intact cerebellar *Ampd3*).

Having confirmed forebrain specific deletion of AMPD, we next characterized its effects on survival, body and brain weight. As expected by the preserved AMPD activity in most body regions, the lifespan of cdKO mice, as well as the body weight, was comparable to their control littermates (Fig S3C). We only detected a mild reduction of brain weight as mice aged (Fig S3C), which suggested age-dependent progression of neurodegeneration. To validate this observation, we analyzed brain sections histologically. Like p15-20 dKO mice, 5-week-old cdKO mice exhibited thinner hippocampal CA1-3 and DG and pyknotic cells in DG (Fig 5A). Remarkably, only the DG region of the cdKO mice showed a severe tissue loss at 8^th^ week of age (Fig 5B), while the CA1-3 regions remained similar to controls over time (Fig 5B and S4D).

**Figure 5:**
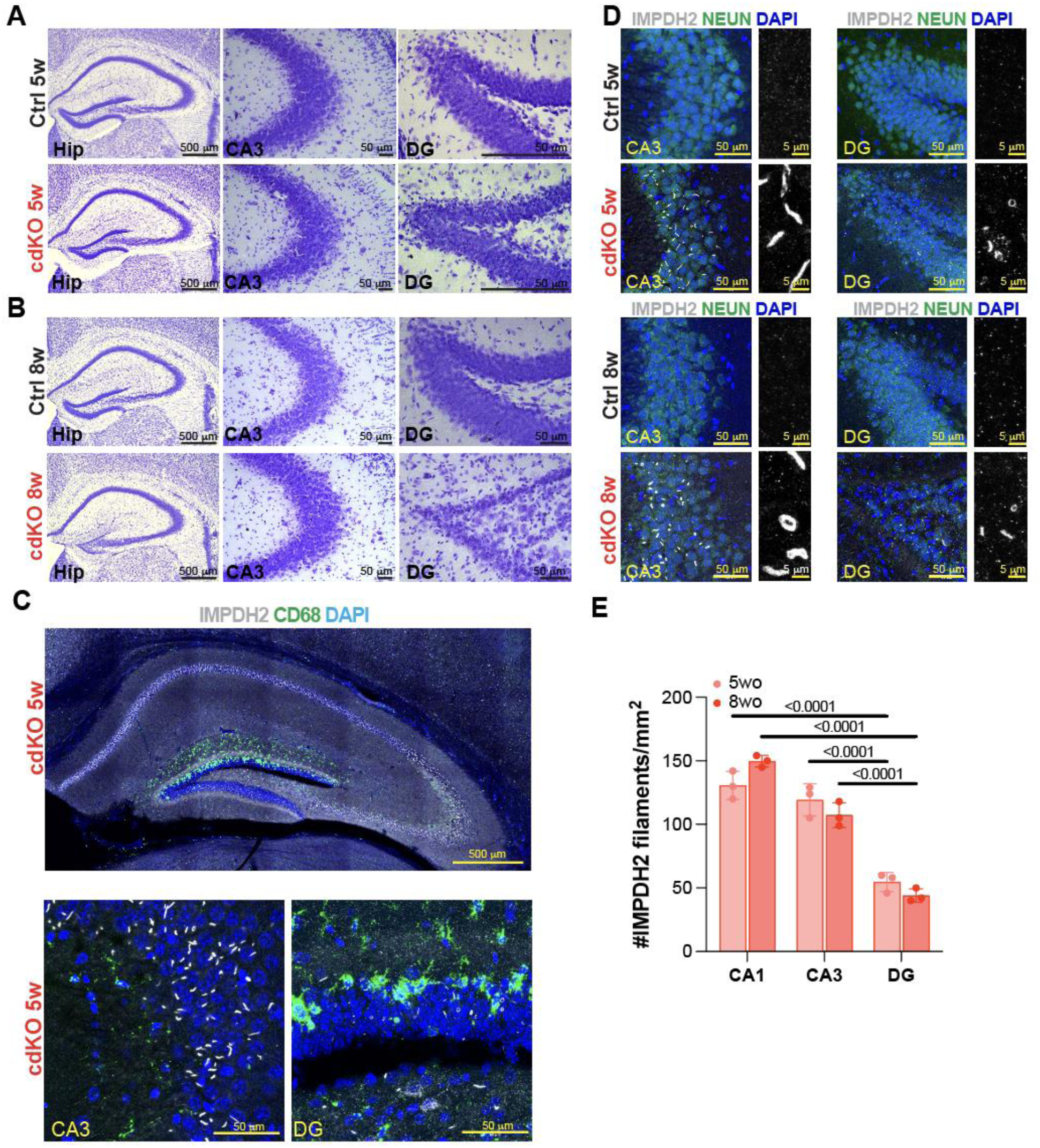
IMPDH2 filaments protect hippocampal CA3 region from neurodegeneration in cerebral cortex and hippocampus specific *Ampd2* and *Ampd3* knock out mice (cdKO) **(A)** Representative Nissl staining of brain sections showing full hippocampus and CA3 and DG regions in 5-week-old (5w) control (Ctrl) and conditional double knockout mice (cdKO). Pyknotic cells are present in cdKO DG region. **(B)** Representative Nissl staining of brain sections showing full hippocampus and CA3 and DG regions in 8-week-old (8w) Ctrl and cdKO mice. Images highlight thinner DG region in cdKO compared to Ctrl hippocampus. **(C)** Representative immunostainings of IMPDH2 and CD68 in cdKO mouse hippocampus. Bottom left panel shows magnification of CA3 region highlighting absence of CD68+ microglia and high density of IMPDH2 filaments. Bottom right panel shows magnification of DG region highlighting presence of CD68+ microglia and barely detectable IMPDH2 filaments. **(D)** Representative immunostainings of IMPDH2 and NEUN in brain sections show persistent presence of IMPDH2 filaments in cdKO CA3 region and low density of small IMPDH2 filaments in cdKO DG. Thinner cdKO DG at 5w and 8w is consistent with data from Nissl staining. **(E)** Bar graph shows mean ± SD of IMDPH2 filaments density observed in different hippocampal regions (CA1, CA3 and DG) in n=3 cDKO mice at 5w and 8w. Significance was calculated with one way ANOVA and Sidak’s post hoc analysis for multiple comparison. Hip=Hippocampus; CA3=Cornu Ammonis; DG=Dentate Gyrus.

We next assessed whether DG degeneration in cdKO mice was associated with a selective microglia accumulation. Consistent with our findings in dKO mice, CD68+ microglia accumulation was predominant in hippocampal DG region of cdKO mice, while CA1-3 regions were only sparsely labeled with CD68+, including at older ages (Fig 5C and S3D). Furthermore, IMPDH2 immunofluorescence stainings showed low density and small IMPDH2 filaments in cdKO DG region at the onset of neurodegeneration (5 weeks of age) and after (8 weeks of age) (Fig 5D-E). In contrast, microglia free CA1-3 regions persistently accumulated large and dense IMPDH2 filaments (Fig 5D-E). Together, this data suggests that IMPDH2 filament accumulation protects from neurodegeneration and microglia infiltration in AMPD deficient mice hippocampus.

### Protection by IMPDH2 filaments is conserved in human AMPD2 deficient patient derived neural cells

Our previous work uncovered that AMPD2 deficient PCH9 patients’ neural progenitor cells (NPCs) are vulnerable to adenosine supplementation in culture (Akizu *et al*., 2013). However, at standard culture conditions PCH9 patients’ NPCs grow as control NPCs, suggesting compensatory mechanisms that support cell growth despite AMPD2 deficiency (Akizu *et al*., 2013). To test whether IMPDH2 filament assembly could be involved in the compensatory mechanism, we derived NPCs from a PCH9 patient induced pluripotent stem cell (iPSCs) and a healthy control family member. PCH9 NPCs showed expression of PAX6 and NESTIN neural progenitor markers, and cell growth comparable to control NPCs (Fig 6A) under standard media conditions. Furthermore, as we predicted, anti-IMPDH2 immunostaining revealed IMPDH2 filaments in PCH9 NPCs but not in controls (Fig 6B) demonstrating that IMPDH2 filament assembly is conserved in PCH9 NPCs.

**Figure 6:**
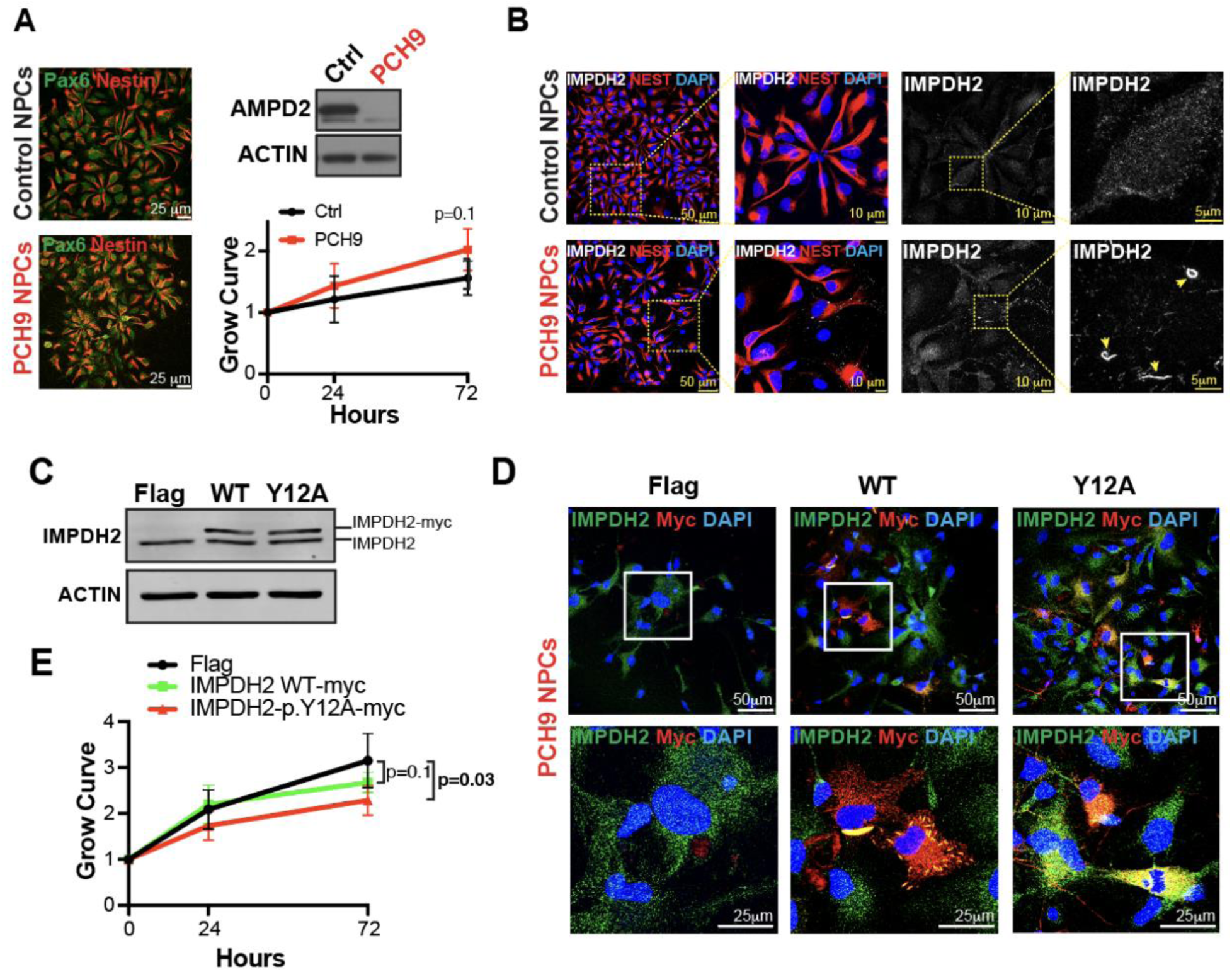
IMPDH2 filaments protect PCH9 patient’s NPCs from growth restrictions. **(A)** Representative images of PCH9 patient and unaffected control NPCs immunostained with anti-PAX6 and anti-NESTIN NPC markers. Western Blot analysis showing lack of AMPD2 protein in PCH9 NPCs compared to controls. Graph shows growth curves of control (Ctrl) and PCH9 NPCs over 72h period. Data shows mean ± SD of 4 independent cultures. Statistical difference is calculated comparing slops of simple linear regression. **(B)** Representative images of anti-IMPDH2 and anti-NESTIN immunostainings showing IMPDH2 filaments in PCH9 NPCs but not in controls. Yellow insets show IMPDH2 filament structures in PCH9 NPCs. **(C)** Western blot of PCH9 NPCs transduced with FLAG peptide (as control), IMPDH2 WT-myc (WT) or a non-polymerizing but catalytically active IMPDH2p.Y12A-myc variant that acts as a dominant negative for filament polymerization. **(C)** Immunostaining of IMPDH2 and MYC in PCH9 NPCs transduced with FLAG, IMPDH2WT-myc (WT) or IMPDH2p.Y12A-myc showing absence of filaments in IMPDH2p.Y12A-myc expressing cells. **(D)** Growth curves quantified with MTT assay show slower growth in IMPDH2p.Y12A-myc transduced NPCs than in FLAG control. Data shows mean ± SD of 6 independent cultures. Statistical difference is calculated comparing slops of simple linear regression.

We next tested the effect of IMPDH2 filaments in cell growth and survival. To this end, we took advantage of a non-polymerizing but catalytically active IMPDH2-p.Y12A variant previously identified(Duong-Ly *et al*., 2018). Overexpression of this variant has a dominant negative effect on IMPDH2 polymerization, preventing endogenous IMPDH2 filament assembly while maintaining the catalytic activity(Duong-Ly *et al*., 2018). Thus, we transduced PCH9 NPCs with lentiviruses carrying either wild type (WT) or p.Y12A IMPDH2, both tagged with a MYC peptide. By WB analysis we confirmed that IMPDH2 WT and IMPDH2 p.Y12A were overexpressed at similar levels (Fig 6C). Furthermore, as previously reported in other cell lines(Anthony *et al*, 2017), we confirmed that IMPDH2 WT overexpression in NPCs induces IMPDH2 filament polymerization while IMPDH2-Y12A expressing NPCs lack IMPDH2 filaments (Fig 6D). Finally, we assessed the effect of IMPDH2 p.Y12A overexpression in PCH9 NPCs growth during a period of 72h and compared with PCH9 NPCs expressing a FLAG peptide or IMPDH2 WT as controls. Results showed no significant differences between IMPDH2 WT and Flag overexpressing PCH9 NPCs (Fig 6D). However, the IMPDH2 p.Y12A overexpression reduced the growth of PCH9 NPCs (Fig 6E). Overall, these data indicate that exogenously inducing IMPDH2 polymerization does not provide a growth advantage, likely due to limiting availability of GTP precursors (i.e. IMP). However, inhibiting IMPDH2 filament assembly by IMPDH2-p.Y12A overexpression is detrimental for PCH9 NPCs growth, suggesting that IMPDH2 polymerization provides a growth advantage to AMPD deficient neural cells.

## Discussion

Selective neuronal vulnerability to metabolic dysfunction is a common feature of pediatric neurodegenerative disorders, but mechanisms that confer vulnerability or neuroprotection are largely unknown. Here we show that IMPDH2 polymerization into micron-sized filaments provides resistance to neuronal death in models of PCH9, a pediatric neurodegenerative disorder caused by AMPD2 deficiency. Remarkably, upregulation of IMPDH2, which often triggers its polymerization, is reported in Alzheimer’s diseases postmortem tissue and mouse models (Neuner *et al*, 2017; Puthiyedth *et al*, 2016; Xu *et al*, 2019), suggesting a broader implication of IMPDH2 polymerization in neurodegeneration. Therefore, our work rises the possibility for a role of IMPDH2 polymerization in neuroprotection and treatment of neurodegenerative disorders.

IMPDH2 catalyzes the rate limiting step for the biosynthesis of guanine nucleotides from IMP. Its activity is tightly regulated at multiple levels, including transcriptionally, posttranscriptionally, and allosterically by binding to purine nucleotides (Hedstrom, 2009). Cellular conditions that deprive guanine nucleotide levels induce IMPDH2 upregulation and conformational changes that boost enzymatic activity, which are reversed once GTP levels are restored (Hedstrom, 2009). Furthermore, IMPDH2 polymerization into micron-sized filaments is emerging as a mechanism that reduces the sensitivity of IMPDH2 to feedback inhibition by guanine nucleotides (Fernandez-Justel *et al*, 2019; Johnson & Kollman, 2020). Therefore, under conditions of high GTP demand, IMPDH2 assembly into filaments would enhance GTP synthesis above the homeostatic levels (Johnson & Kollman, 2020). Accordingly, IMPDH2 filaments have been reported in highly proliferative cells, such as pluripotent stem cells (Carcamo *et al*, 2011; Keppeke *et al*, 2018) and activated T-lymphocytes (Calise *et al*, 2018; Duong-Ly *et al*., 2018), and upon pharmacological treatments that block IMP to GTP flux in cell cultures (Ji *et al*., 2006; Thomas *et al*, 2012). Our work is the first showing IMPDH2 filaments in postmitotic neurons and in pathological conditions. Remarkably, we show that only few neurons exposed to AMPD deficiency and GTP depletion assemble IMPDH2 into filaments suggesting that IMPDH2 polymerization occurs under specific intracellular conditions. Whether these conditions depend on cell specific energy supply, expression of specific proteins or posttranslational modifications of IMPDH2, remain to be determined. However, our work suggests that IMP availability relative to GTP (i.e. IMP/GTP ratio) may be involved in the induction of IMPDH2 polymerization. This is consistent with previous work showing that increasing concentrations of IMP and ATP induce IMPDH2 polymerization *in vitro* (Johnson & Kollman, 2020). In agreement, we also uncovered that IMPDH2 filaments are in close proximity with mitochondria, a primary source of ATP that may favor local conditions for IMPDH2 polymerization. These observations, suggest that neurons in the hippocampal dentate gyrus, where IMPDH2 filaments are scarce, may have lower IMP and ATP availability, making them more vulnerable to succumb to GTP deprivation than neurons in the CA1-3 hippocampal regions, which are rich in IMPDH2 filaments. Although, current limitations for single cell resolution of metabolomic methods preclude us from testing this hypothesis, the model that emerges from our study is that neurons with available guanine nucleotide precursors can survive to AMPD deficiency associated nucleotide imbalances through IMPDH2 filament assembly to boost GTP synthesis.

Our work is also consistent with species-specific differences in brain purine nucleotide requirements. Indeed, while hippocampal dentate gyrus degenerates in AMPD deficient mice, there is currently no data showing hippocampal degeneration in human AMPD2 deficiency. Furthermore, the cerebellum, which is hypoplastic at birth in human patients, remains apparently intact in AMPD deficient mice, at least up to their death before postnatal day 21. Much of the cerebellar pathology in human AMPD2 deficiency arises during development (Kortum *et al*., 2018) and, remarkably, there are marked differences in human and mouse cerebellar development, including the timing and progenitor cell types (Haldipur *et al*, 2019). These differences may be responsible of disparities between human and mouse cerebellar vulnerability to AMPD deficiency. However, it is also possible that the cerebellum would degenerate if AMPD deficient mice survived past day 21. Indeed, lack of IMPDH2 filaments in the cerebellum predicts its susceptibility to degenerate in AMPD deficient mice. Thus, depleting AMPD in cerebella, like we did for the forebrain in dKO mice, may provide a mouse model to longitudinally study cerebellar requirements for purine nucleotides and IMPDH2 filament assembly in the future.

Whether IMPDH2 and its polymerization into filaments confers resistance to other neurodegenerative conditions is question that remains to be determined. Interestingly, emerging data also suggest this possibility. Indeed, recent studies implicate *de novo IMPDH2* mutations in the etiology of neurodevelopmental disorders (Kuukasjarvi *et al*, 2021; O’Neill *et al*., 2023; Zech *et al*, 2020). Although most of the disease variants impair the sensitivity of IMPDH2 for GTP mediated inhibition, one of these variants (p.S160del) leads to inability of the protein to assemble into filaments (O’Neill *et al*., 2023). Therefore, it will be interesting to determine if the patient carrying this variant develops a neurodegenerative condition over time. In contrast, mutations in the retinal paralogue, IMPDH1, which are a major cause of photoreceptor degenerative disorders, suggest the opposite. Some of these mutations impair the disassembly of IMPDH1 filaments upon guanine nucleotide supplementation *in vitro.* Consequently, these mutant IMDPH1 variants constitutively polymerize when overexpressed in cells (Fernandez-Justel *et al*., 2019; Keppeke *et al*, 2023). Furthermore, recent data shows that the long-term overexpression of a constitutively polymerizing IMPDH1 disease variant induces cell death in HEp-2 cells line. Consistently, insoluble nuclear inclusions of IMPDH1 have been recently proposed as a mechanism contributing to age related dysfunction of dopaminergic neurons of the substantia nigra (Woulfe *et al*, 2024). These data indicate data constitutive IMPDH1/2 filament formation may be as detrimental as the inability to form filaments and suggest that the coordination between IMPDH1/2 activity demand and its polymerization are critical to support cell survival. In agreement, our work shows that despite supporting PCH9 NPC growth, further induction of IMPDH2 filaments, by IMPDH2WT overexpression, does not improve the cell growth, likely due to lack of enough IMP to be fueled into IMPDH2 filaments for guanine nucleotide biosynthesis in PCH9 NPCs.

Nonetheless, if IMPDH2 filament assembly prevents neurodegeneration, could we leverage this for treatment? Our current work suggests that in a scenario where GTP precursors (i.e. IMP) are available, IMPDH2 filaments support cell growth. Interestingly, we previously found that the treatment of PCH9 patients’ NPCs with 5-Aminoimidazole-4-carboxamide riboside (AICAR) prevents neurodegeneration by increasing de novo IMP synthesis and restoring GTP levels (Akizu *et al*., 2013). Remarkably, evidence shows that AICAR induces IMPDH2 polymerization (Schiavon *et al*, 2018) suggesting it could potentially at as a dual booster rescuing survival of AMPD2 deficient neurons. Furthermore, AICAR enhances exercise endurance and transiently promotes neurogenesis and learning and memory in healthy mice (Guerrieri & van Praag, 2015; Narkar *et al*, 2008). Yet, there is also evidence showing that chronic AICAR supplementation may elicit neuronal apoptosis, and impair axon growth and synaptic plasticity (Guerrieri & van Praag, 2015),(Ma *et al*, 2014; Williams *et al*, 2011). Therefore, although AICAR may not be the best therapeutic candidate, our work opens the possibility for IMPHD2 polymerization combined with *de novo* purine nucleotide stimulation as a therapeutic target for neurodegeneration, at least in PCH9.

## Methods

### Mice

All animal procedures were performed according to NIH guidelines and approved by the Institutional Animal Care and Use Committee (IACUC) at Children’s Hospital of Philadelphia. *Ampd2*-/- and *Ampd3* first knock outs mice were generated previously(Akizu *et al*., 2013; Toyama *et al*., 2012). *Ampd2*-/- and *Ampd3-/-* double knock out (dKO) mice were generated by breeding *Ampd2^-/+^;Ampd3^-/-^* males with females. Given that *Ampd2^-/+^;Ampd3^-/-^* are grossly indistinguishable from wild type mice(Akizu *et al*., 2013), these were used as control littermates for all our experiments. For conditional AMPD deficient mice (cdKO), we first excised the trapping cassette of the *Ampd3^-/-^*first knock out allele by FRT mediated recombination after breeding it into ACTB:FLPe (Jackson Laboratory # 005703) transgenic mice. Mice carrying Ampd3 conditional allele (*Ampd3^f/f^*), with loxp sites flanking exon 3, were then breed into *Ampd2^-/-^* background and *Emx1:Cre* knock in mice (Jackson Laboratory #005628)(Gorski *et al*, 2002). To generate *Ampd2^-/-^*; *Ampd3^f/f^*; *Emx1-Cre^+/-^*mice (cdKO) for experiments, *Ampd2^-/-^*; *Ampd3^f/f^*mice were bred with *Ampd2^+/-^*; *Ampd3^f/f^*; *Emx1-Cre^+/-^* mice. Littermates with *Ampd2^-/-^*; *Ampd3^f/f^* or *Ampd2^+/-^*; *Ampd3^f/f^*; *Emx1-Cre^+/-^* genotypes were used as controls for experiments. Male and female mice were indistinguishably used for experiments. Mice were housed two to five animal per cage with a 12-h light-dark cycle (lights on from 0600 to 1800h) at constant temperature (23°C) with ad libitum access to food and water. Routine genotyping was performed by tail biopsy and PCR as previously described(Akizu *et al*., 2013).

### Human neural progenitor cell culturing

Approval to work with previously reported PCH9 iPSCs(Akizu *et al*., 2013) was obtained by the Children’s Hospital of Philadelphia Institutional Review Board. Neural progenitor cells (NPCs) were generated from PCH9-1236 patient iPSCs cultured on Matrigel (Corning, #354277) coated dishes with mTSER1 (STEM-Cell Technologies #85850) media as previously described(Akizu *et al*., 2013). Briefly, embryoid bodies (EBs) were formed by mechanical dissociation of iPSC colonies and plated in neural induction media (DMEM F12, 1x N2, 1x B27, 1μM LDN and 1μM SB431542) under rotation at 75 rpm for 9 days. Resultant EBs were then plated on Matrigel coated dishes in NBF medium (DMEM F12, 1x N2, 1x B27, 20 ng/ml bFGF, 1x Penicillin-Streptomycin). Neural rosettes were visible to pick after 4-6 days, and NPCs dissociated with Accutase (Thermo Fisher, A1110501) and plated on Matrigel with NBF media. Media was replaced every other day and NPCs passaged once a week approximately. All experiments were performed with NPCs at passage 5-8.

### NPC Growth Curve Assessment

NPCs were plated at 15x10^3^ cells/well in 96 well plates and allowed to adhere overnight. For growth curve analysis, viable number of cells were measured with MTT (Sigma-Aldrich M2128-1G) at 0h (the day after plating), 24h and 72h. Briefly, 0.325 mg/ml MTT (3-(4,5-Dimethylthiazol-2-yl)-2,5-Diphenyltetrazo-lium Bromide; Sigma) was added into culture medium for 2h and incubated at 37C. Media was then aspired and DMSO added to lyse NPCs and dissolve the purple formazan crystals formed by the reduction of yellow MTT in viable cells. Absorbance was measured in SpectraMax 190 Microplate reader at 570 nm and subtracted for background signal at 670 nm.

### Transduction of NPCs

NPCs were transduced with lentiviruses carrying Flag, IMPDH2WT-Myc or IMPDH2p.Y12A-Myc cDNA. To this end, IMPDH2WT-Myc and IMPDH2p.Y12A-Myc in pcDNA3 plasmids(Anthony *et al*., 2017) (a gift from Jeffrey Peterson) were subcloned into pINDUCER20(Meerbrey *et al*, 2011). Lentiviruses were then produced in 10 cm dishes of Hek293T cells, by co-transfection of 10 μg pINDUCER20 plasmids, 5 μg p8.9NdeltaSB (Addgene #132929) and 0.5 μg pCMV-VSV-G (Addgene #8454)) with Lipofectamine 2000 (Invitrogen #11668500). 8h after transfection, media was replaced with NBF media and 48h after media with lentiviruses was transferred to NPCs in the presence of 8 mg/ml polybrene. Following one-week selection with 200 mg/ml G418, NPCs were treated with 100 ng/ml doxycycline for the transgene expression and plated for experiments.

### Immunoblotting

Mice were euthanized with CO_2_ followed by decapitation, and the region of interest (hippocampus, cortex and cerebellum) was rapidly dissected and frozen in liquid nitrogen and stored at -80°C until further processing. Brain and NPC samples were lysed with RIPA buffer (Cell signaling, 9806S-CST) supplemented with protease inhibitors (Sigma, P8340). After 30 min of incubation at 4°C, lysate was centrifuged at 12,000 rpm for 10 min. Protein concentration was determined by Bradford (Thermo Fisher 23246) method according to the manufacturer’s instructions. Protein samples were diluted in an equal volume of 2x LDS sample buffer (Thermo Fisher, B0007) and supplemented with DTT to a final concentration of 50mM (Bio Rad, 1610611). Protein samples (30-50 μg) were separated on 10-18% SDS-PAGE gels and transferred to PVDF membrane, stained with Ponceau S, and blocked with 5% milk in TBS-Tween 20 for 2h and incubated with primary antibodies diluted in the 5% BSA in TBS-Tween 20 overnight at 4°C (AMPD2, Sigma HPA045760, 1:1000; IMPDH2, Abcam ab129165, 1:1000; S6, Santa Cruz sc74459, 1:1000; p-S6, Cell signaling 5364, 1:1500; Tubulin, Sigma T6074, 1:2000; Actin, Gen Script A00702-40, 1:2000). Membranes were washed 3 times with TBS-Tween 20 and incubated with anti-mouse IgG-HRP (Thermo Fisher SA1100; Li-cor Biosciences 926-68072) or anti-rabbit IgG-HRP (Thermo Fisher 31458; Li-cor Biosciences 926-32213) secondary antibodies (all 1:4000) for 2h at room temperature. After washing 3 times with TBS-Tween 20, membranes were exposed to enhanced chemiluminescence substrate (Thermo Fisher 34076) and signal developed in autoradiography films and AFP Mini-Med 90 X-Ray Film Processor. Bands were quantified using ImageJ (NIH, USA) or Image Studio Lite (USA).

### Histology and Immunocytochemistry

Mice were anaesthetized with isoflurane (1-4%) and intracardially perfused with PBS followed by 4% PFA in PBS. Brains were extracted and further fixed in 4% PFA overnight at 4°C and washed 3 times with PBS. Coronal or sagittal sections of 50 μm were obtained in a vibratome, mounted on slides and dried overnight at room temperature. Dried sections were stained with cresyl violet (Millipore C5042) followed by dehydration in alcohol gradient and clearing with xylene before mounting with Permount (Thermo Fisher SP15) mounting media. Stained brain sections were imaged with an epifluorescent microscope (Leica DM6000).

For immunofluorescence staining, 50 μm thick brain sections were permeabilized and blocked with blocking solution (5% fetal serum bovine plus 0.5% Triton X-100 in PBS) for 2h at room temperature. Sections were incubated with primary antibodies (IMPDH2, Abcam ab129165, 1:1000; CD68, Bio Rad MCA1957, 1:250; NEUN, Millipore MAB377, 1:2000; GFAP, Aves Lab 75-240, 1:1000, S6, Santa Cruz sc74459, 1:1000; p-S6, Cell signaling 5364, 1:1500) in blocking solution overnight at 4°C. Section were washed 3 times with 0.1% Triton X100 in PBS and incubated with secondary antibodies (Alexa Fluor 488-, 555-633-labelled goat anti-mouse, goat anti-rabbit, goat anti-rat or anti-chicken IgGs (H+L) 1:500, Thermo Fisher) and DAPI (1:100, Thermo Fisher, D3571) in blocking solution at room temperature for 2h. Then, sections were washed 3 times with 0.1% Triton X100 in PBS and mounted in coverslips Prolong Gold mounting media. Imaging was performed using a Leica SP8 confocal microscope. For z-stack images, 5μm z-stack confocal images were acquired at 1μm intervals. Image processing was performed using ImageJ Software (NIH, USA).

### Electron microscopy

Mice were anaesthetized with isoflurane (1-4%) and intracardially perfused with 2%PFA+2% Glutaraldehyde in 0.1M sodium cacodylate, followed by post-fixation in 2% osmium tetroxide, dehydrated through a graded ethanol series, infiltrated in propylene oxide and embedded in resin. Semithin sections were stained with toluidine blue and ultrathin sections were stained with lead citrate. Images were captured with a Zeiss Libra I20 TEM.

### RNA sequencing and data analysis

20 day-old mice were euthanized and cortex, hippocampus and cerebellum were dissected on ice, fast frozen, and stored in -80°C until RNA extraction. On the day of RNA extraction, 50-100mg tissue from each sample was lysed in 1ml TRIzol (Invitrogen, #15596026) according to the manufacturer’s recommendations. RNA integrity and quantity were assessed in Bioanalyzer. Strand-specific mRNA-seq libraries were generated at Novogene and sequenced on an Illumina NovaSeq 6000 platform with 2x150 PE configuration at an average of 15 million reads per sample. Adapter and poor quality sequences were trimmed, and cleaned mRNA reads mapped to the *Mus musculus* GRCm38 reference genome using STAR (v2.7.3a)(Dobin *et al*, 2013). Reads were counted using FeatureCounts from the subread package (v2.0.1)(Liao *et al*, 2014) with parameters: -p -C -O. TPM values were calculated from featureCounts-derived counts by normalizing the count rates to gene lengths and then scaling them by the sum of all normalized count rates, multiplied by 106. Heatmap of gene expression was generated using the tidyverse R package with t.test for statistical comparison between genotypes. Differential gene expression analysis was performed with DEseq2 (V1.38.3)(Love *et al*, 2014) excluding genes with less than five reads, and all genes from chromosome X and Y. Differential expression was performed using a linear model and as factor the genotypes. Raw p-values were adjusted using the Benjamini-Hochberg method. Differentially expressed genes were defined as having an adjusted p value of less than 0.05. Volcano plots of differentially expressed genes were generated with the EnhancedVolcano R package. Gene Set Enrichment Analysis (GSEA) was performed on the cell-type specific gene lists from the top 500 significantly enriched genes in Microglia_1, Microglia_2, Microglia_3, DAM_Microglia and Perivascular_MF (renamed as Macrophage) extracted from(Keren-Shaul *et al*., 2017). GSEA was performed with the clusterProfiler(Wu *et al*, 2021) R package with raw p-values adjusted with the Benjamini-Hochberg method. GSEA-heatmap was generated with the help of the tidyverse R package. Functional annotation enrichment analysis was performed using Enrichr(Chen *et al*, 2013) and sub-category MSigDB Hallmark 2000. ENRICHR false discovery rate (FDR) values were calculated using the Benjamini–Hochberg test in ENRICHR and P values were calculated using Fisher’s exact test in ENRICHR. Significantly enriched (pAdj < 0.05 (-log10=1.301) with Benjamini–Hochberg correction) categories were represented as bubble plots generated in GraphPad Prism v.10.

### Nucleotide analysis by LC-MS

Mice were intraperitoneally injected with pentobarbital (100mg/Kg) and hippocampus, cortex and cerebellum were harvested and placed immediately in liquid nitrogen. Frozen brain regions were lyophilized and then homogenized in 80% methanol using a Precellys homogenizer. Aliquots of homogenates were treated with additional methanol and then centrifugated to precipitate proteins. The supernatants were dried under nitrogen and the resuspended in HPLC mobile phases for resolution in an Agilent 1290 HPLC coupled with an Agilent triple quadruple 6495B mass spectrometer (Agilent, Wilmington, DE). Extracted samples were separated on an Agilent ZORBAX Eclipse Plus (2.1x100mm, 1.8 um) at 50°C using 9.5 min linear gradient from 99% solvent A (0.1% formic acid in water) to 100% solvent B (acetonitrile with 0.1% formic acid) at flow rate of 0.8ml/min. The gradient used was as following: 2.5 min, 100%A, 0%B; 3.5min, 99.2%A, 0.8%B; 5.0min, 98.2%A, 3.2%B; 7.0min, 0%A, 100%B; 7.5 min, 0%A, 100%B; 7.6min, 100%A, 0%B. The MS was operated in the positive ion mode using electrospray ionization. The source gas temperature was 250°C, the gas flow was 11L/min, and the nebulizer was set at 50 psi. The amount of each nucleotide was determined by comparison to standards with known concentration.

### Statistical analysis

All experiments were performed in at least three independent biological replicates. The n number for each experiment, details of statistical analysis and software are described in the figure legends or main text. Statistical analysis was performed using Prism (GraphPad).

## Acknowledgments

We thank Shannon Modla at Delaware University Biotechnology Institute electron microscopy core for processing and imaging tissue for transmission electron microscopy and the Penn Metabolomics Core (RRID:SCR_022381) in the Cardiovascular Institute at the University of Pennsylvania for nucleotide metabolomics analyses. We are also grateful to Jeffrey R. Peterson, who inspired this work and shared insightful knowledge in addition to pCDNA3 plasmids, and especially, for inspiring this work with his insights on IMPDH2 filaments and passion for science. This work was supported by the National Institute of Health NIH/NINDS R00NS089859 grant (N.A.) and International Brain Research Organization ISN-Research Postdoctoral Fellowship (M.F-M.).

## Author contribution

Study conceptualization and design: M.F-M. and N.A. Maintenance of mouse colony: M.F-M., L.O., K.W. and N.A. Experimental execution, and data collection: M.F-M., L.O., Y.Z., J.A.T-H., K.W. and N.A. Computational analyses: T.R. Data interpretation: M.F-M., X.O-G. N.A. Manuscript preparation: M.F-M. and N.A. Manuscript edit and review: All authors

## Conflict of Interest statement

None declared.

## Data availability

RNAseq data was deposited in GEO under the GSE253045 (https://www.ncbi.nlm.nih.gov/geo/query/acc.cgi?acc=GSE253045) accession number. All the other data are available in the main text or the supplementary materials.

---

**Figure S1:**
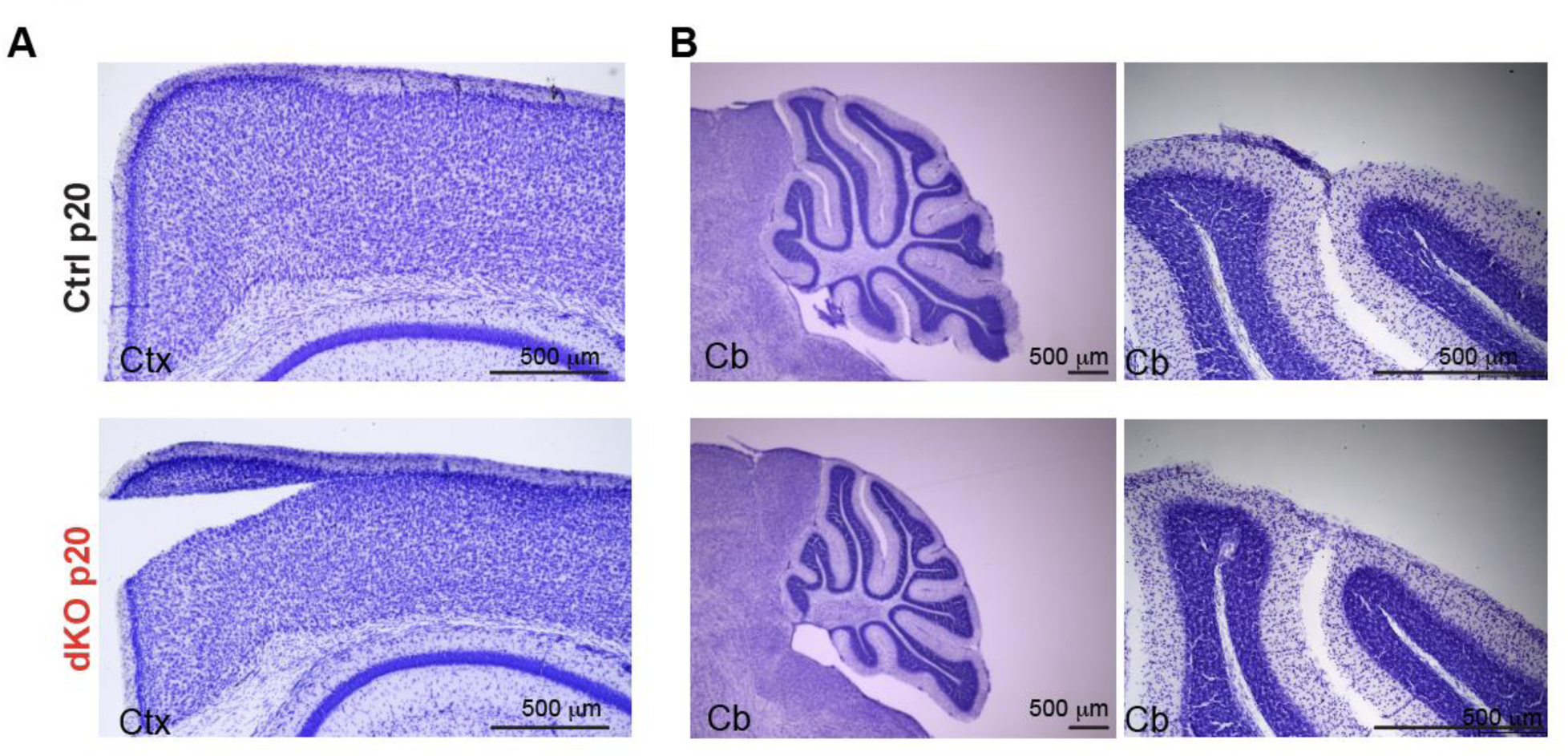
No signs of overt neurodegeneration in cortex and cerebellum of Ampd2 and Ampd3 double knockout (dKO) mice. **(A-B)** Representative image of the cerebral cortex (A) and cerebellum (B) of Control (Ctrl) and double knockout (dKO) mice brain vibratome sections at postnatal day 20 (p20) showing intact tissue, with no structural abnormalities or no signs of neurodegeneration. Ctx=Cortex and Cb=Cerebellum.

**Figure S2:**
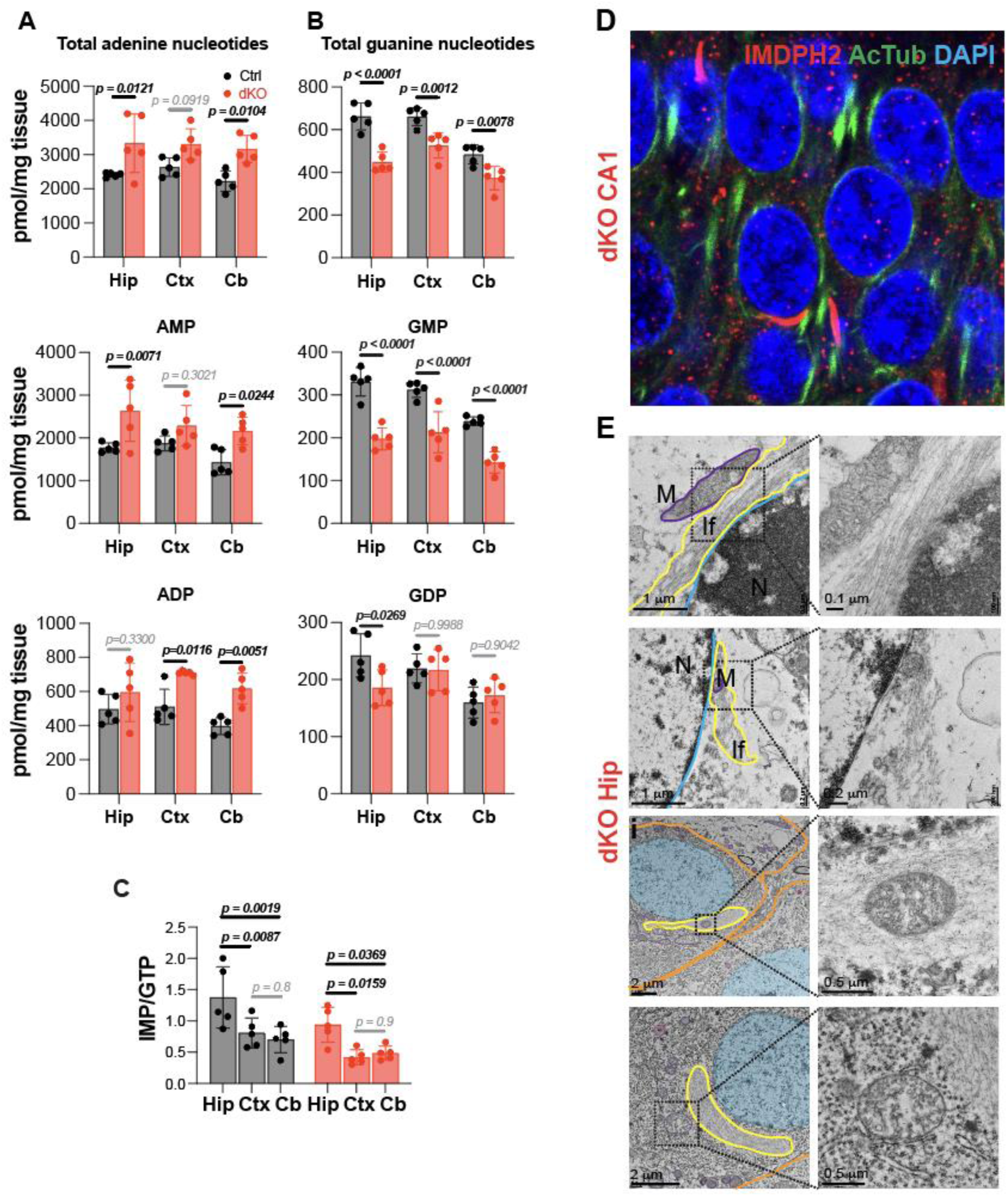
Purine nucleotide imbalance and IMPDH2 filament assembly in dKO mice brain regions. **(A-B)** Adenine (A) and Guanine (B) nucleotides levels in Hippocampus (Hip), Cortex (Ctx) and Cerebellum (Cb) of control (Ctrl) and double knockout (dKO) mice at postnatal day 20 (p20). Graph shows the mean ± SD of n=5 mice per genotype. Significance was calculated with one way ANOVA with Sidak’s post hoc analysis for multiple comparison**. (C)** Ratio between IMP and GTP levels is larger in control and dKO mice (p20) hippocampus than in Cortex and Cerebellum. Graph shows mean ± SD of n=5 mice per genotype. Significance was calculated with one way ANOVA and Sidak’s post hoc analysis for multiple comparison**. (D)** Immunostaining of IMPDH2 and Acetylated tubulin (primary cilia marker) in dKO mouse at p20. **(E)** Representative transmission electron microscopy images show IMPDH2 filaments (IF) (traced in yellow) within hippocampal neurons. Magnifications (left panels) show IMPDH2 filaments close to mitochondria (M) (traced in purple) and nucleus (N) (blue).

**Figure S3:**
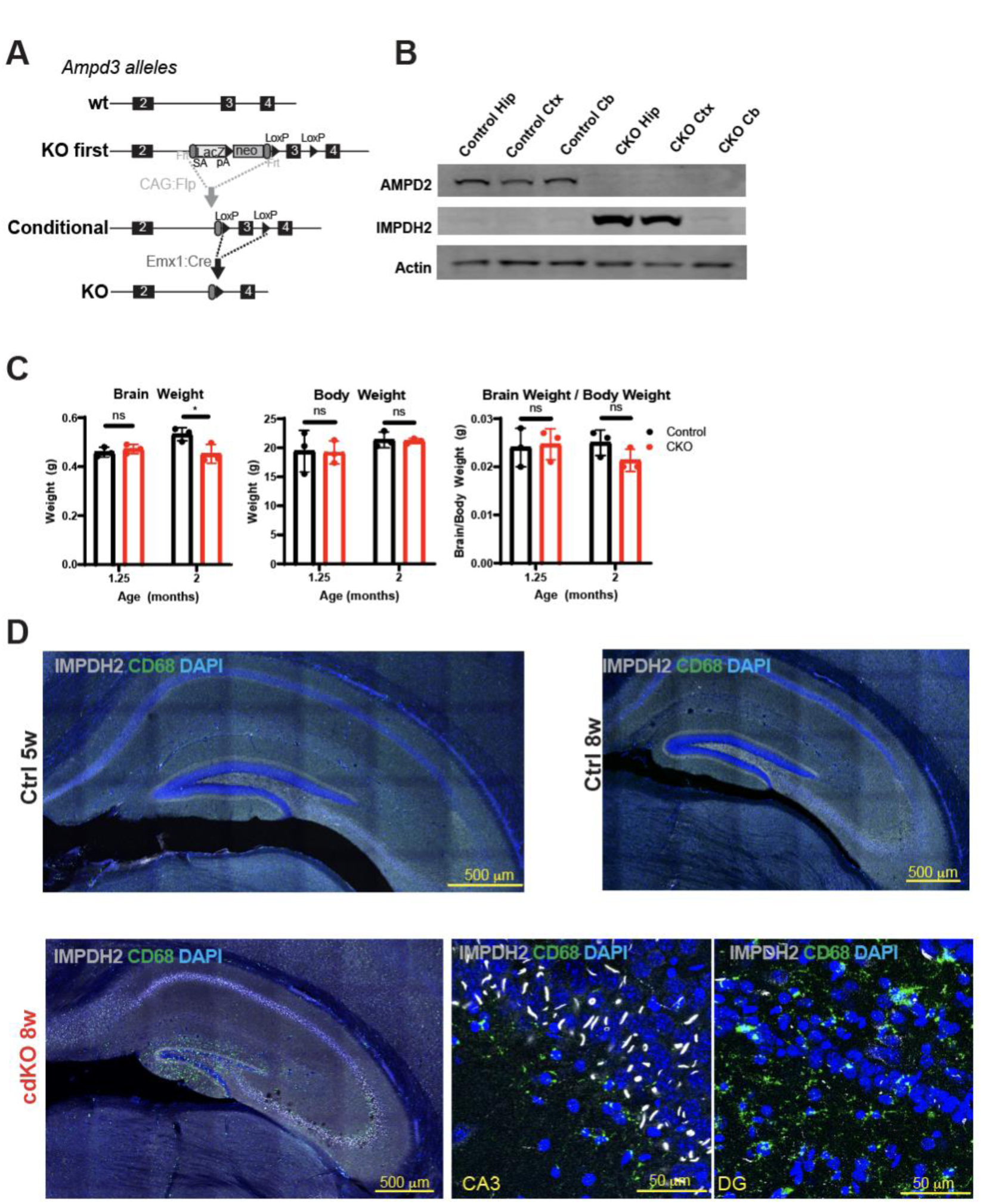
Forebrain deletion of *Ampd2* and *Ampd3* leads to mild loss of brain weight, hippocampal DG accumulation of microglia and CA3 accumulation of IMPDH2 filaments. (A) Schematic diagram showing Cre-mediated excision of floxed Ampd3 exon 3 to generate the conditional double knockout (cdKO) mouse for *Ampd2* and *Ampd3*. **(B)** Representative western blot showing absence of AMPD2 and upregulation of IMPDH2 in cdKO compared with controls mice (5w). **(C)** Brain weight, body weight and their ratio at different age. Graph shows mean ± SD of n= 3 mice per genotype. Significance was calculated with two sided unpaired t-test**. (D)** Top images are representative immunostainings of IMPDH2 and CD68 in control brain sections (at 5 and 8 weeks of age) showing absence of IMPDH2 filaments and microglia in control mice. Bottom panels show representative images of cdKO hippocampus immunostained for IMPHD2 and CD68. Magnification of CA3 region image shows high density of IMPDH2 filaments and low density of microglia. Magnification of DG region shows neurons absence of IMPDH2 filaments and CD68+ microglia.

## Notes

### Competing Interest Statement

The authors have declared no competing interest.

## References

1. Akizu N, Cantagrel V, Schroth J, Cai N, Vaux K, McCloskey D, Naviaux RK, Van Vleet J, Fenstermaker AG, Silhavy JL et al (2013) AMPD2 regulates GTP synthesis and is mutated in a potentially treatable neurodegenerative brainstem disorder. Cell 154: 505–517

2. An S, Kumar R, Sheets ED, Benkovic SJ (2008) Reversible compartmentalization of de novo purine biosynthetic complexes in living cells. Science 320: 103–106

3. Anthony SA, Burrell AL, Johnson MC, Duong-Ly KC, Kuo YM, Simonet JC, Michener P, Andrews A, Kollman JM, Peterson JR (2017) Reconstituted IMPDH polymers accommodate both catalytically active and inactive conformations. Mol Biol Cell 28: 2600–2608

4. Arts WF, Loonen MC, Sengers RC, Slooff JL (1993) X-linked ataxia, weakness, deafness, and loss of vision in early childhood with a fatal course. Ann Neurol 33: 535–539

5. Badimon A, Strasburger HJ, Ayata P, Chen X, Nair A, Ikegami A, Hwang P, Chan AT, Graves SM, Uweru JO et al (2020) Negative feedback control of neuronal activity by microglia. Nature 586: 417–423

6. Becker MA, Puig JG, Mateos FA, Jimenez ML, Kim M, Simmonds HA (1988) Inherited superactivity of phosphoribosylpyrophosphate synthetase: association of uric acid overproduction and sensorineural deafness. Am J Med 85: 383–390

7. Bowne SJ, Sullivan LS, Blanton SH, Cepko CL, Blackshaw S, Birch DG, Hughbanks-Wheaton D, Heckenlively JR, Daiger SP (2002) Mutations in the inosine monophosphate dehydrogenase 1 gene (IMPDH1) cause the RP10 form of autosomal dominant retinitis pigmentosa. Hum Mol Genet 11: 559–568

8. Buey RM, Ledesma-Amaro R, Velazquez-Campoy A, Balsera M, Chagoyen M, de Pereda JM, Revuelta JL (2015) Guanine nucleotide binding to the Bateman domain mediates the allosteric inhibition of eukaryotic IMP dehydrogenases. Nat Commun 6: 8923

9. Calise SJ, Abboud G, Kasahara H, Morel L, Chan EKL (2018) Immune Response-Dependent Assembly of IMP Dehydrogenase Filaments. Front Immunol 9: 2789

10. Camici M, Micheli V, Ipata PL, Tozzi MG (2010) Pediatric neurological syndromes and inborn errors of purine metabolism. Neurochem Int 56: 367–378

11. Carcamo WC, Satoh M, Kasahara H, Terada N, Hamazaki T, Chan JY, Yao B, Tamayo S, Covini G, von Muhlen CA et al (2011) Induction of cytoplasmic rods and rings structures by inhibition of the CTP and GTP synthetic pathway in mammalian cells. PLoS One 6: e29690

12. Chen EY, Tan CM, Kou Y, Duan Q, Wang Z, Meirelles GV, Clark NR, Ma’ayan A (2013) Enrichr: interactive and collaborative HTML5 gene list enrichment analysis tool. BMC Bioinformatics 14: 128

13. de Brouwer AP, Williams KL, Duley JA, van Kuilenburg AB, Nabuurs SB, Egmont-Petersen M, Lugtenberg D, Zoetekouw L, Banning MJ, Roeffen M et al (2007) Arts syndrome is caused by loss-of-function mutations in PRPS1. Am J Hum Genet 81: 507–518

14. Dobin A, Davis CA, Schlesinger F, Drenkow J, Zaleski C, Jha S, Batut P, Chaisson M, Gingeras TR (2013) STAR: ultrafast universal RNA-seq aligner. Bioinformatics 29: 15–21

15. Duong-Ly KC, Kuo YM, Johnson MC, Cote JM, Kollman JM, Soboloff J, Rall GF, Andrews AJ, Peterson JR (2018) T cell activation triggers reversible inosine-5’-monophosphate dehydrogenase assembly. J Cell Sci 131

16. Escobar-Henriques M, Daignan-Fornier B (2001) Transcriptional regulation of the yeast gmp synthesis pathway by its end products. J Biol Chem 276: 1523–1530

17. Fernandez-Justel D, Nunez R, Martin-Benito J, Jimeno D, Gonzalez-Lopez A, Soriano EM, Revuelta JL, Buey RM (2019) A Nucleotide-Dependent Conformational Switch Controls the Polymerization of Human IMP Dehydrogenases to Modulate their Catalytic Activity. J Mol Biol 431: 956–969

18. Fishbein WN, Armbrustmacher VW, Griffin JL (1978) Myoadenylate deaminase deficiency: a new disease of muscle. Science 200: 545–548

19. French JB, Jones SA, Deng H, Pedley AM, Kim D, Chan CY, Hu H, Pugh RJ, Zhao H, Zhang Y et al (2016) Spatial colocalization and functional link of purinosomes with mitochondria. Science 351: 733–737

20. Fu R, Sutcliffe D, Zhao H, Huang X, Schretlen DJ, Benkovic S, Jinnah HA (2015) Clinical severity in Lesch-Nyhan disease: the role of residual enzyme and compensatory pathways. Mol Genet Metab 114: 55–61

21. Gorski JA, Talley T, Qiu M, Puelles L, Rubenstein JL, Jones KR (2002) Cortical excitatory neurons and glia, but not GABAergic neurons, are produced in the Emx1-expressing lineage. J Neurosci 22: 6309–6314

22. Guerrieri D, van Praag H (2015) Exercise-mimetic AICAR transiently benefits brain function. Oncotarget 6: 18293–18313

23. Gunter JH, Thomas EC, Lengefeld N, Kruger SJ, Worton L, Gardiner EM, Jones A, Barnett NL, Whitehead JP (2008) Characterisation of inosine monophosphate dehydrogenase expression during retinal development: differences between variants and isoforms. Int J Biochem Cell Biol 40: 1716–1728

24. Haldipur P, Aldinger KA, Bernardo S, Deng M, Timms AE, Overman LM, Winter C, Lisgo SN, Razavi F, Silvestri E et al (2019) Spatiotemporal expansion of primary progenitor zones in the developing human cerebellum. Science 366: 454–460

25. Hartman SC, Buchanan JM (1959) Nucleic acids, purines, pyrimidines (nucleotide synthesis). Annu Rev Biochem 28: 365–410

26. Hedstrom L (2009) IMP dehydrogenase: structure, mechanism, and inhibition. Chem Rev 109: 2903–2928

27. Hoxhaj G, Hughes-Hallett J, Timson RC, Ilagan E, Yuan M, Asara JM, Ben-Sahra I, Manning BD (2017) The mTORC1 Signaling Network Senses Changes in Cellular Purine Nucleotide Levels. Cell Rep 21: 1331–1346

28. Ji Y, Gu J, Makhov AM, Griffith JD, Mitchell BS (2006) Regulation of the interaction of inosine monophosphate dehydrogenase with mycophenolic Acid by GTP. J Biol Chem 281: 206–212

29. Johnson MC, Kollman JM (2020) Cryo-EM structures demonstrate human IMPDH2 filament assembly tunes allosteric regulation. Elife 9

30. Juda P, Smigova J, Kovacik L, Bartova E, Raska I (2014) Ultrastructure of cytoplasmic and nuclear inosine-5’-monophosphate dehydrogenase 2 “rods and rings” inclusions. J Histochem Cytochem 62: 739–750

31. Kennan A, Aherne A, Palfi A, Humphries M, McKee A, Stitt A, Simpson DA, Demtroder K, Orntoft T, Ayuso C et al (2002) Identification of an IMPDH1 mutation in autosomal dominant retinitis pigmentosa (RP10) revealed following comparative microarray analysis of transcripts derived from retinas of wild-type and Rho(-/-) mice. Hum Mol Genet 11: 547–557

32. Keppeke GD, Chang CC, Peng M, Chen LY, Lin WC, Pai LM, Andrade LEC, Sung LY, Liu JL (2018) IMP/GTP balance modulates cytoophidium assembly and IMPDH activity. Cell Div 13: 5

33. Keppeke GD, Chang CC, Zhang Z, Liu JL (2023) Effect on cell survival and cytoophidium assembly of the adRP-10-related IMPDH1 missense mutation Asp226Asn. Front Cell Dev Biol 11: 1234592

34. Keren-Shaul H, Spinrad A, Weiner A, Matcovitch-Natan O, Dvir-Szternfeld R, Ulland TK, David E, Baruch K, Lara-Astaiso D, Toth B et al (2017) A Unique Microglia Type Associated with Restricting Development of Alzheimer’s Disease. Cell 169: 1276–1290 e1217

35. Kim HJ, Sohn KM, Shy ME, Krajewski KM, Hwang M, Park JH, Jang SY, Won HH, Choi BO, Hong SH et al (2007) Mutations in PRPS1, which encodes the phosphoribosyl pyrophosphate synthetase enzyme critical for nucleotide biosynthesis, cause hereditary peripheral neuropathy with hearing loss and optic neuropathy (cmtx5). Am J Hum Genet 81: 552–558

36. Kortum F, Jamra RA, Alawi M, Berry SA, Borck G, Helbig KL, Tang S, Huhle D, Korenke GC, Hebbar M et al (2018) Clinical and genetic spectrum of AMPD2-related pontocerebellar hypoplasia type 9. Eur J Hum Genet 26: 695–708

37. Kuukasjarvi A, Landoni JC, Kaukonen J, Juhakoski M, Auranen M, Torkkeli T, Velagapudi V, Suomalainen A (2021) IMPDH2: a new gene associated with dominant juvenile-onset dystonia-tremor disorder. Eur J Hum Genet 29: 1833–1837

38. Lane AN, Fan TW (2015) Regulation of mammalian nucleotide metabolism and biosynthesis. Nucleic Acids Res 43: 2466–2485

39. Lesch M, Nyhan WL (1964) A Familial Disorder of Uric Acid Metabolism and Central Nervous System Function. Am J Med 36: 561–570

40. Liao LX, Song XM, Wang LC, Lv HN, Chen JF, Liu D, Fu G, Zhao MB, Jiang Y, Zeng KW et al (2017) Highly selective inhibition of IMPDH2 provides the basis of antineuroinflammation therapy. Proc Natl Acad Sci U S A 114: E5986–E5994

41. Liao Y, Smyth GK, Shi W (2014) featureCounts: an efficient general purpose program for assigning sequence reads to genomic features. Bioinformatics 30: 923–930

42. Love MI, Huber W, Anders S (2014) Moderated estimation of fold change and dispersion for RNA-seq data with DESeq2. Genome Biol 15: 550

43. Ma T, Chen Y, Vingtdeux V, Zhao H, Viollet B, Marambaud P, Klann E (2014) Inhibition of AMP-activated protein kinase signaling alleviates impairments in hippocampal synaptic plasticity induced by amyloid beta. J Neurosci 34: 12230–12238

44. Meerbrey KL, Hu G, Kessler JD, Roarty K, Li MZ, Fang JE, Herschkowitz JI, Burrows AE, Ciccia A, Sun T et al (2011) The pINDUCER lentiviral toolkit for inducible RNA interference in vitro and in vivo. Proc Natl Acad Sci U S A 108: 3665–3670

45. Narkar VA, Downes M, Yu RT, Embler E, Wang YX, Banayo E, Mihaylova MM, Nelson MC, Zou Y, Juguilon H et al (2008) AMPK and PPARdelta agonists are exercise mimetics. Cell 134: 405–415

46. Neuner SM, Wilmott LA, Hoffmann BR, Mozhui K, Kaczorowski CC (2017) Hippocampal proteomics defines pathways associated with memory decline and resilience in normal aging and Alzheimer’s disease mouse models. Behav Brain Res 322: 288–298

47. Novarino G, Fenstermaker AG, Zaki MS, Hofree M, Silhavy JL, Heiberg AD, Abdellateef M, Rosti B, Scott E, Mansour L et al (2014) Exome sequencing links corticospinal motor neuron disease to common neurodegenerative disorders. Science 343: 506–511

48. O’Neill AG, Burrell AL, Zech M, Elpeleg O, Harel T, Edvardson S, Mor-Shaked H, Rippert AL, Nomakuchi T, Izumi K et al (2023) Neurodevelopmental disorder mutations in the purine biosynthetic enzyme IMPDH2 disrupt its allosteric regulation. J Biol Chem 299: 105012

49. Pareek V, Tian H, Winograd N, Benkovic SJ (2020) Metabolomics and mass spectrometry imaging reveal channeled de novo purine synthesis in cells. Science 368: 283–290

50. Pascual O, Casper KB, Kubera C, Zhang J, Revilla-Sanchez R, Sul JY, Takano H, Moss SJ, McCarthy K, Haydon PG (2005) Astrocytic purinergic signaling coordinates synaptic networks. Science 310: 113–116

51. Pierre G (2013) Neurodegenerative disorders and metabolic disease. Arch Dis Child 98: 618–624

52. Puthiyedth N, Riveros C, Berretta R, Moscato P (2016) Identification of Differentially Expressed Genes through Integrated Study of Alzheimer’s Disease Affected Brain Regions. PLoS One 11: e0152342

53. Saudubray JM, Garcia-Cazorla A (2018) An overview of inborn errors of metabolism affecting the brain: from neurodevelopment to neurodegenerative disorders. Dialogues Clin Neurosci 20: 301–325

54. Schiavon CR, Griffin ME, Pirozzi M, Parashuraman R, Zhou W, Jinnah HA, Reines D, Kahn RA (2018) Compositional complexity of rods and rings. Mol Biol Cell 29: 2303–2316

55. Sperling O, Eilam G, Sara Persky B, De Vries A (1972) Accelerated erythrocyte 5-phosphoribosyl-1-pyrophosphate synthesis. A familial abnormality associated with excessive uric acid production and gout. Biochem Med 6: 310–316

56. Synofzik M, Muller vom Hagen J, Haack TB, Wilhelm C, Lindig T, Beck-Wodl S, Nabuurs SB, van Kuilenburg AB, de Brouwer AP, Schols L (2014) X-linked Charcot-Marie-Tooth disease, Arts syndrome, and prelingual non-syndromic deafness form a disease continuum: evidence from a family with a novel PRPS1 mutation. Orphanet J Rare Dis 9: 24

57. Taylor JP, Hardy J, Fischbeck KH (2002) Toxic proteins in neurodegenerative disease. Science 296: 1991–1995

58. Thomas EC, Gunter JH, Webster JA, Schieber NL, Oorschot V, Parton RG, Whitehead JP (2012) Different characteristics and nucleotide binding properties of inosine monophosphate dehydrogenase (IMPDH) isoforms. PLoS One 7: e51096

59. Toyama K, Morisaki H, Cheng J, Kawachi H, Shimizu F, Ikawa M, Okabe M, Morisaki T (2012) Proteinuria in AMPD2-deficient mice. Genes Cells 17: 28–38

60. van Dijk T, Baas F, Barth PG, Poll-The BT (2018) What’s new in pontocerebellar hypoplasia? An update on genes and subtypes. Orphanet J Rare Dis 13: 92

61. Watts RW (1983) Some regulatory and integrative aspects of purine nucleotide biosynthesis and its control: an overview. Adv Enzyme Regul 21: 33–51

62. Williams T, Courchet J, Viollet B, Brenman JE, Polleux F (2011) AMP-activated protein kinase (AMPK) activity is not required for neuronal development but regulates axogenesis during metabolic stress. Proc Natl Acad Sci U S A 108: 5849–5854

63. Wong V (1997) Neurodegenerative diseases in children. Hong Kong Med J 3: 89–95

64. Woulfe J, Munoz DG, Gray DA, Jinnah HA, Ivanova A (2024) Inosine monophosphate dehydrogenase intranuclear inclusions are markers of aging and neuronal stress in the human substantia nigra. Neurobiol Aging 134: 43–56

65. Wu T, Hu E, Xu S, Chen M, Guo P, Dai Z, Feng T, Zhou L, Tang W, Zhan L et al (2021) clusterProfiler 4.0: A universal enrichment tool for interpreting omics data. Innovation (Camb) 2: 100141

66. Xu J, Patassini S, Rustogi N, Riba-Garcia I, Hale BD, Phillips AM, Waldvogel H, Haines R, Bradbury P, Stevens A et al (2019) Regional protein expression in human Alzheimer’s brain correlates with disease severity. Commun Biol 2: 43

67. Zakaria RBM, Malta M, Pelletier F, Addour-Boudrahem N, Pinchefsky E, Martin CS, Srour M (2023) Classic “PCH” Genes are a Rare Cause of Radiologic Pontocerebellar Hypoplasia. Cerebellum

68. Zech M, Jech R, Boesch S, Skorvanek M, Weber S, Wagner M, Zhao C, Jochim A, Necpal J, Dincer Y et al (2020) Monogenic variants in dystonia: an exome-wide sequencing study. Lancet Neurol 19: 908–918

69. Zimmermann AG, Gu JJ, Laliberte J, Mitchell BS (1998) Inosine-5’-monophosphate dehydrogenase: regulation of expression and role in cellular proliferation and T lymphocyte activation. Prog Nucleic Acid Res Mol Biol 61: 181–209

70. Zydowo MM, Purzycka-Preis J, Ogasawara N (1989) Deficiency of AMP deaminase in human erythrocytes. Adv Exp Med Biol 253A: 31–34

